# Pathophysiological regulation of lung function by the free fatty acid receptor FFA4

**DOI:** 10.1101/2020.05.18.101170

**Authors:** Rudi Prihandoko, Davinder Kaur, Coen H. Wiegman, Elisa Alvarez-Curto, Chantal Donovan, Latifa Chachi, Trond Ulven, Martha R. Tyas, Eloise Euston, Zhaoyang Dong, Abdulrahman Ghali M Alharbi, Richard Kim, Jack G. Lowe, Philip M. Hansbro, Kian Fan Chung, Christopher E. Brightling, Graeme Milligan, Andrew B. Tobin

## Abstract

Increased prevalence of inflammatory airway diseases including asthma and chronic obstructive pulmonary disease (COPD) together with a significant number of patients being inadequately controlled by current frontline treatments means that there is a need to define novel therapeutic targets for these conditions^1^. Here we investigate a member of the G protein-coupled receptor (GPCR) family, FFA4, which responds to free circulating fatty acids, including dietary omega-3 fatty acids found in fish oils^2–4^. Although usually associated with metabolic responses linked with food intake, we show that FFA4 is expressed in the lung where it is coupled to G_q/11_-signalling. Activation of FFA4 by drug-like agonists produced relaxation of murine airway smooth muscle mediated, at least in part, by the release of the prostaglandin PGE_2_ that subsequently acts on EP_2_ prostanoid receptors. In normal mice, activation of FFA4 resulted in a decrease in lung resistance. Importantly, in acute and chronic ozone models of pollution-mediated inflammation, and in house-dust mite and cigarette smoke-induced inflammatory disease, FFA4 agonists acted to reduce airway resistance, whilst this response was absent in mice lacking expression of FFA4. The expression profile of FFA4 in human lung was very similar to that observed in mice and the response to FFA4/FFA1 agonists similarly mediated human airway smooth muscle relaxation. Hence, our study provides evidence that pharmacological targeting of lung FFA4, and possibly combined activation of FFA4 and FFA1, has *in vivo* efficacy that might have therapeutic value in the treatment of bronchoconstriction associated with inflammatory airway diseases such as asthma and COPD.

## Background

Asthma and chronic obstructive pulmonary disease (COPD) are chronic respiratory inflammatory diseases that affect >500 million people worldwide, causing substantial morbidity and exacting a considerable cost to health services^5–7^. Despite the effectiveness of current therapies the heterogeneity of these conditions means that ∼45% of asthmatics remain uncontrolled and in COPD the combination of corticosteroid, long-acting muscarinic antagonists and *β*-adrenoceptor agonists inadequately control symptoms and exacerbations in many patients^5-8^. We have addressed the urgent need to understand novel paradigms in lung physiology that might reveal new therapeutic strategies by focusing on the role and potential clinical value of a member of the G protein-coupled receptor (GPCR) super-family, FFA4 (previously known as GPR120). This GPCR responds to a range of free circulating long chain (>C12) fatty acids (LCFAs) that include the diet-derived essential fatty acids linoleic and *α*-linolenic acid as well as omega-3 polyunsaturated fatty acids prevalent in fish oils^2–4^. The majority of previous studies on FFA4 have focused on the regulation of physiological responses associated with food intake. Expressed on entero-endocrine cells of the gut^9, 10^, various cell types in pancreatic islets^11, 12^, white adipose tissue^13^, and on immune cells, particularly macrophages^14^, FFA4 has been strongly implicated in the regulation of glucose homeostasis^12^ and inflammation, particularly in connection with adipose tissue in animals fed on a high fat diet^15^. These data, together with the emergence of a number of small *“drug-like”* ligands to FFA4^3^, has fuelled interest from both academia and industry in targeting FFA4 for the treatment of metabolic disease including type II diabetes and obesity^4^.

It has however recently become clear that FFA4 is abundantly expressed in lung epithelial cells^16^ and a recent study suggested that omega-3 fatty acids acting via FFA4 receptors might have some beneficial effects on airway epithelial repair following naphthalene-induced airway injury^17^. Other than this study, the role that FFA4 might play in lung physiology is completely unknown. Here, we use a combination of novel pharmacological agents, genetically engineered mice and both *ex vivo* and *in vivo* techniques, to determine that FFA4 receptors expressed in mouse and human airways, mediate airway smooth muscle relaxation in a manner that is relevant in both normal physiology and inflammatory airway disease.

### Expression of FFA4 in mouse airways

Using both RT-PCR (**fig 1a**) and qRT-PCR (**fig 1b**) we confirmed the presence of FFA4 mRNA in the mouse lung at levels substantially higher than seen for a second receptor for LCFAs, FFA1 (**figs 1a,b**) and confirmed the lack of expression of FFA4 mRNA in equivalent samples from a genetically engineered mouse strain where the first exon of the FFA4 gene was replaced with the coding sequence of *β*-galactosidase (FFA4-KO(*β*gal) mice) (**figs 1a,b**). To investigate the tissue distribution of FFA4 further, we stained for *β*-galactosidase in FFA4-KO(*β*gal) mice as this acts as a surrogate marker for FFA4 expressing cells. Using this approach we established that FFA4 is expressed primarily in cells of the airway epithelium (**fig 1c**). These data are supported by studies using an in-house generated mouse FFA4 selective antiserum^18^ where it was confirmed that FFA4 expression was primarily associated with the epithelium of the mid- and lower-airways, with very much lower levels of expression associated with airway smooth muscle (ASM), as defined by the presence of *α*-actin (**fig 1d**). Co-staining with an antiserum to club cell specific protein 10 (CC10) demonstrated that at least one cell type expressing FFA4 in the airway epithelium was club cells (**fig 1e**), a result consistent with a recent mRNA analysis that identified high level enrichment of FFA4 transcript levels in club cells^19^.

**Fig 1.**
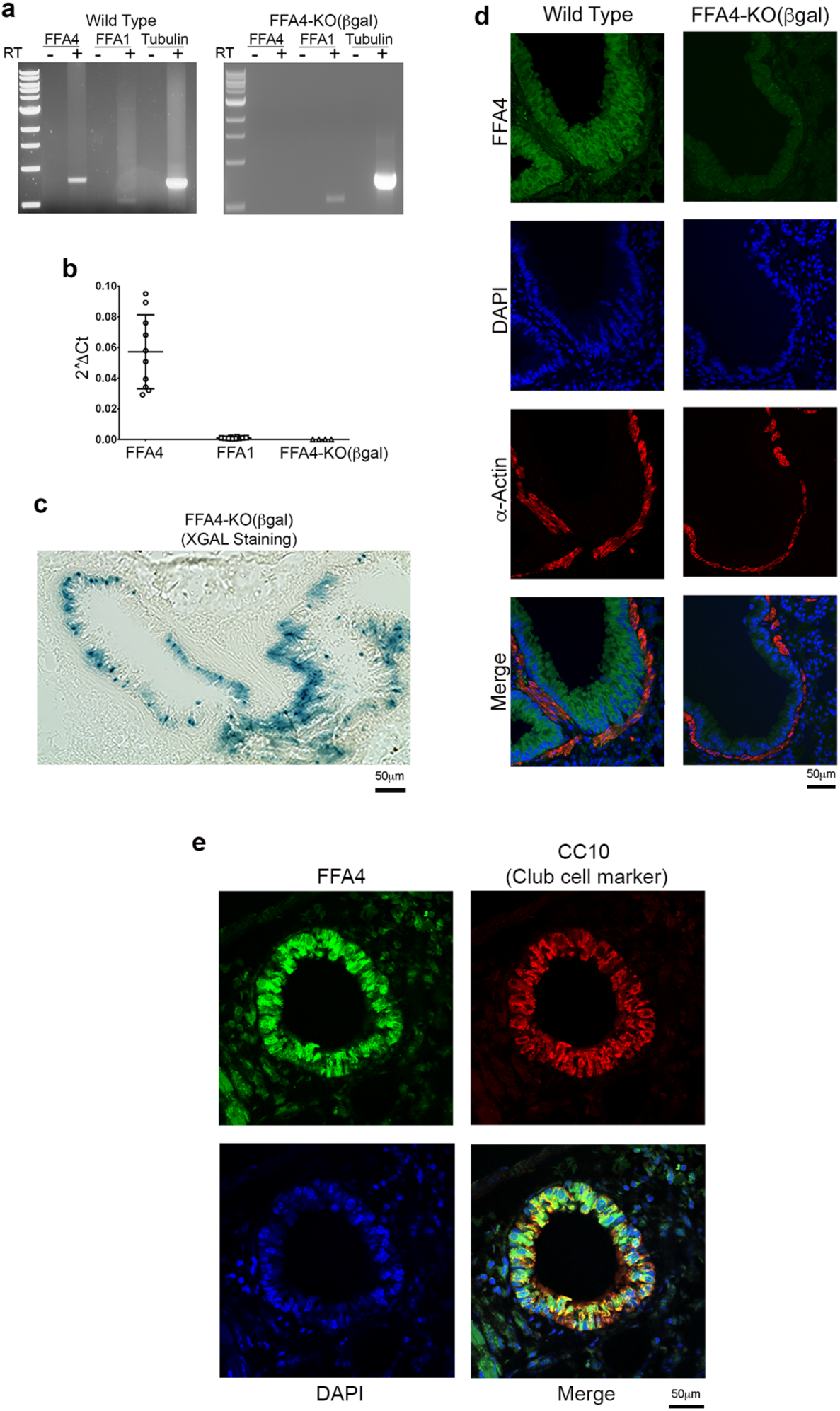
FFA4 receptors are expressed in murine lung and localise primarily to the epithelial layer. **(a)** Transcript levels of FFA1, FFA4 and tubulin (as a control) were identified in lung from wild type and FFA4-KO(βgal) mice using RT-PCR. Shown is a representative gel of 3 independent experiments. The expected sizes of PCR products are as follows; FFA4 = 800 base pairs (bp), tubulin = 750 bp and FFA1 = 530 bp. **(b)** Quantitative RT-PCR of the lung tissue samples from wild type and FFA4-KO(βgal) mice was conducted using GAPDH housekeeping gene as a control. Data represent mean ± S.D. of 4-10 independent experiments. **(c)** Representative confocal image of lung tissue sections obtained from FFA4-KO(βgal) mice and stained for β-galactosidase as a surrogate for FFA4 expression. **(d)** Representative images of lung tissue sections obtained from wild-type (left) and FFA4-KO(βgal) (right) mice co-stained with an in-house generated mouse FFA4 specific antiserum (green), α-actin antibody to stain ASM (red) and DAPI to identify nuclei (blue). **(e)** Immunofluorescence co-staining of mouse FFA4 and club cell marker (CC10). The images are representative of at least three independent experiments.

### FFA4 mediates airway smooth muscle relaxation and bronchodilation

To probe the activity of FFA4 we used a well characterized agonist, TUG-891^20–24^, which has activity at both FFA1 and FFA4 (**fig 2a**). In addition we employed the recently generated agonist TUG-1197^25^ that we confirm here selectively activates FFA4 but not FFA1 (**fig 2b**). Importantly, FFA4 has previously been shown to signal in a bimodal fashion, initiating signal transduction via heterotrimeric G proteins as well as through a mechanism operating via the recruitment of arrestin adaptor proteins^18, 20–22^. Our previous studies have established that the recruitment of arrestin3 in response to TUG-891 is nearly totally dependent on receptor phosphorylation and mediates processes such as receptor internalization^18, 21^. Here we show that TUG-1197 acts as an agonist at FFA4 in both G protein-dependent signalling, as measured by G_q/11_-mediated calcium mobilization (**fig 2b**) and activation of Extracellular Regulated Protein Kinase 1/2, (ERK1/2; **fig 2c,d**). In addition, TUG-1197 also is an agonist at promoting arrestin3 recruitment (**fig 2e**), receptor phosphorylation (**fig 2f**) and receptor internalization (**fig 2g**). Hence, the signalling properties of TUG-1197 indicated that this ligand is a selective agonist at FFA4, initiating signalling through canonical G-protein and arrestin pathways. TUG-1197 is therefore a useful tool ligand for probing the physiological activity of FFA4. Moreover, whilst also able to activate FFA1 (**fig 2a**) TUG-891 was able to mimic all the effects of TUG-1197 at FFA4 (**fig 2c-g**).

**Fig 2.**
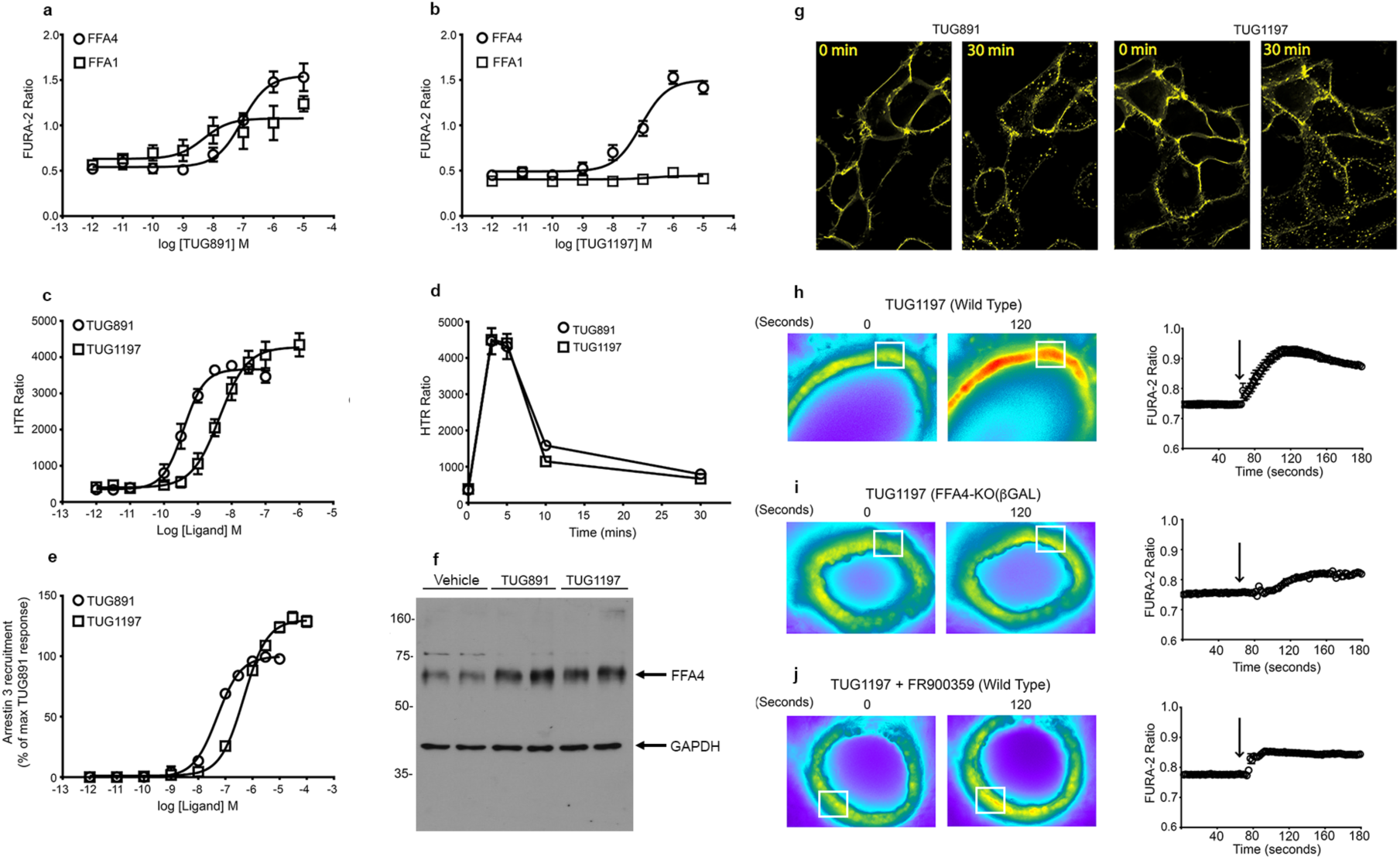
Pharmacology of FFA4 agonist ligands in cell based assays and *ex vivo* lung tissue preparations. **(a,b)** Calcium concentration-response curves in Flp-In T-Rex 293 cells expressing mouse FFA1 or FFA4 stimulated with **(a)** TUG-891 or **(b)** TUG1197. **(c)** ERK1/2 concentration-response curves of CHO Flp-In cells expressing mouse FFA4 following 5 min stimulation with TUG-891 or TUG-1197. **(d)** Kinetics of TUG-891 and TUG-1197 mediated ERK1/2 responses at maximal concentration (1 μM) of each ligand. **(e)** Arrestin3 interaction concentration response curves of Flp-In T-REx 293 cells expressing recombinant mouse FFA4 stimulated with varying concentrations of TUG-891 or TUG-1197. (**f**) FFA4 phosphorylation in response to TUG-891 and TUG-1197 was determined in Western blots of lysates prepared from Flp-In T-REx 293 expressing mouse FFA4 and probed with an anti-phospho antiserum to pThr347/pSer350 on mouse FFA4. Parallel immunoblotting to detect GAPDH acted as a loading control. (**g**) Internalization of eYFP-tagged mouse FFA4 expressed in Flp-In T-REx 293 cells following 30 minute application of TUG-891 or TUG-1197 revealed punctate intracellular localization of FFA4 indicative of internalized receptor. **(h-j)** Elevation of intracellular calcium levels evoked by TUG-1197 in precision cut lung slices from **(h)** wild-type mice, (**i**) FFA4 KO(βgal) mice and **(j)** wild type mice in the presence of the G_q_ inhibitor FR900359. The white box indicates the area of interest where the fluorescence changes (changes in 340/380nm emission from the FURA2-AM calcium indicator (FURA-2 Ratio)) are illustrated in the graph to the right of the images. Arrows indicate the time point at which the FFA4 agonist was added. The experiments shown are representative of at least three independent experiments. (Data in **a-e** are the mean ± S.E.M of at least three independent experiments).

In precision cut lung slices (PCLS) derived from wild type mice, TUG-1197 induced a rapid increase in intracellular calcium (**fig 2h**). In contrast, no significant calcium response to TUG-1197 was observed in PCLS derived from FFA4-KO(*β*gal) mice (**fig 2i**). Furthermore, the calcium response to TUG-1197 in PCLS from wild type mice was significantly inhibited by the G_q/11_-blocker, FR900359^26^ (**fig 2j**). From these data it was concluded that FFA4 in the lung is functionally active and couples to the G_q/11_/phospholipase C/inositol phosphate signal transduction cascade and that the activity of our tool compounds TUG-891 and TUG-1197 in driving a calcium response in lung tissue is exclusively via FFA4.

Since FFA4 mediated a strong calcium response in PCLS the possibility that FFA4 might promote ASM contraction was examined. Application of the bronchoconstrictors, carbachol or serotonin, resulted in narrowing of airways in PCLS **(fig S1a-c**), responses that act here as a positive control. In contrast, the application of TUG-891 or TUG-1197, at concentrations that resulted in robust calcium responses in PCLS (e.g. 50μM (**fig 2h**)) did not result in any significant change in the diameter of airways in PCLS (**fig S1b, c**). These data indicated that FFA4 did not mediate ASM contraction. Instead, experiments designed to test the possibility that FFA4 agonists could promote ASM relaxation revealed that both TUG-891 and TUG-1197 acted in a concentration-dependent manner to increase the diameter of airways pre-contracted with carbachol (**fig 3a-d**) with pEC_50_ values of 4.60 ± 0.10 and 4.77 ± 0.15 respectively. In the case of FFA4-mediated ASM relaxation the maximal effective concentration of either TUG-891 or TUG-1197 returned airways to 70-80% of the diameter observed before carbachol pre-treatment (**fig 3a-d**). Importantly, the response to both TUG-891 and TUG-1197 was significantly reduced by the FFA4 selective antagonist AH7614^27^ (**fig 3e,f and fig S2a,b**). Similarly, ASM relaxation to TUG-1197 was significantly diminished in PCLS derived from FFA4-KO(*β*gal) mice (**fig 3g,h**). In both the antagonist experiments and in FFA4-KO(*β*gal) mice, there was a small response to TUG-1197 and TUG-891 which may represent some off-target effect of these ligands. However, further support for a broncho-relaxation response of FFA4 activation was evident from our studies using a chemically distinct, and previously described, selective FFA4 agonist, Merck Compound A^15^. This agent showed similar FFA4 agonist properties (pEC_50_ and E_max_) to TUG-1197 and TUG-891 in *in vitro* calcium mobilization assays (pEC_50_ values of 6.76, 6.77, 6.00 for TUG-891, TUG-1197 and Compound A, respectively) **(fig S3a).** Importantly, in PCLS, Compound A mimicked the effects of TUG-891 and TUG-1197 in reducing ASM contraction **(fig S3a-c).** Finally, the relaxation response to FFA4 agonists was not restricted to airways pre-treated with carbachol since airways pre-contracted with serotonin (5HT) were also seen to be similarly relaxed in response to TUG-1197 (**fig S4a,b**).

**Fig 3.**
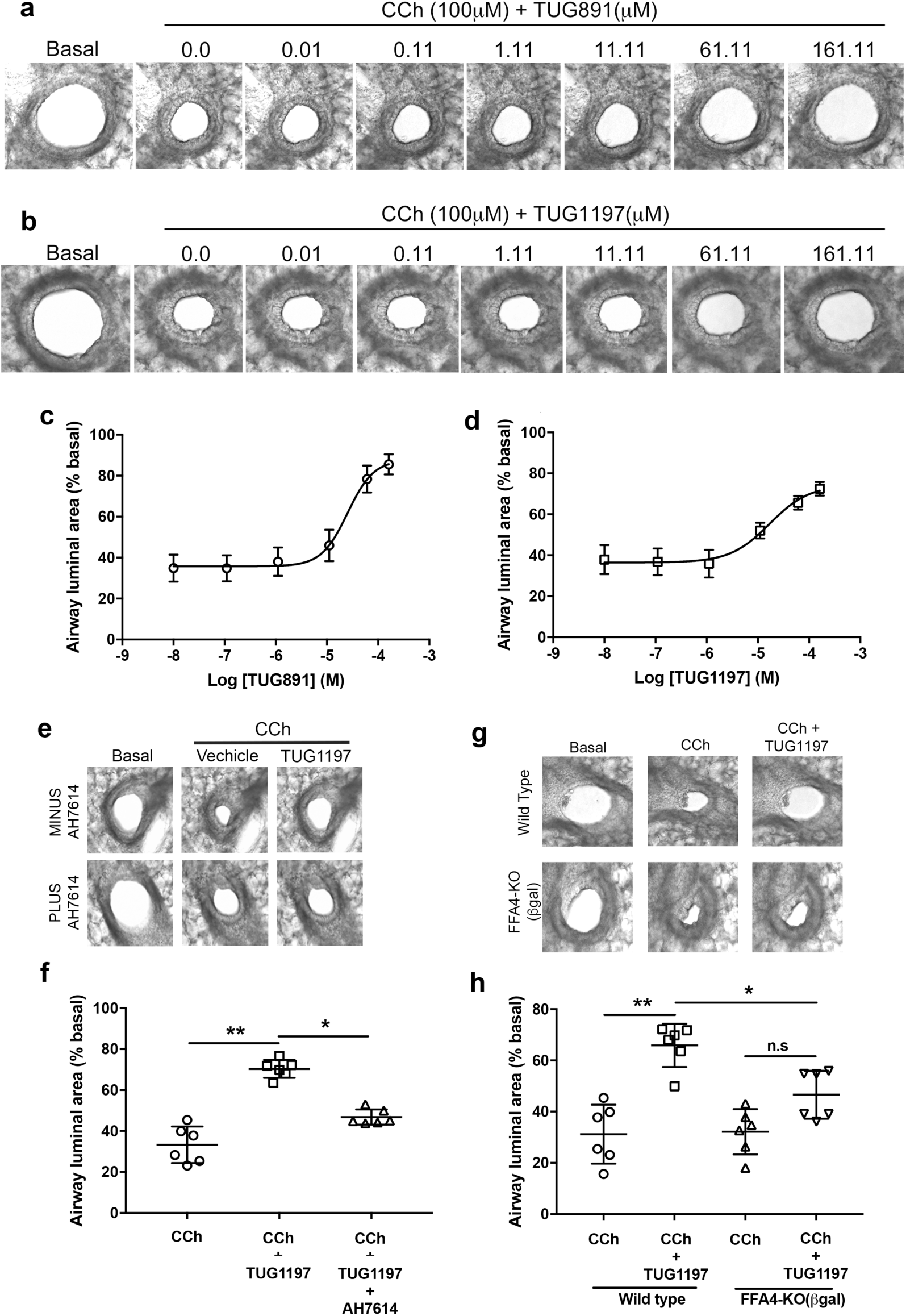
Activation of FFA4 leads to airway relaxation in pre-contracted airways. **(a,b)** Representative images of the concentration-dependent relaxation responses to **(a)** TUG-891 and **(b)** TUG-1197 in precision cut lung slices (PCLS) pre-contracted with carbachol (CCh). **(c,d)** Quantification of the data represented in **a** and **b**. The data presented are the mean ± S.E.M (n=5). **(e)** Representative images of the effect of the FFA4 antagonist AH7614 on the TUG-1197-mediated relaxation response. **(f)** Quantification of the experiment in **e** from at least 5 mice. **(g)** Representative images of the TUG-1197-mediated relaxation responses in PCLS from wild type and FFA4-KO(βgal) mice. (**h**) Quantification of the experiment in **g** from at least 5 mice Data presented in **f** and **h** are the means ± S.E.M. of at least 5 animals where *p<0.05, **p<0.01 as determined by ANOVA with Bonferroni post-hoc test.

### FFA4 mediates bronchodilation in normal lung physiology and disease

The data from the above *ex vivo* experiments led to the prediction that FFA4 agonists would increase the diameter of airways and thereby decrease airway resistance *in vivo*. This was tested in anaesthetised mice where airway resistance was elevated via administration of nebulized acetylcholine^28^. Under these conditions, the response to acetylcholine alone in FFA4-KO(*β*gal) was comparable to that observed in wild-type mice (fig 4a (compare vehicle treated mice). The effect of activation of FFA4 on the acetylcholine response was tested by co-administration of nebulized TUG-891 or TUG-1197 with acetylcholine. Under these conditions the acetylcholine-mediated increase in resistance was significantly attenuated (**fig 4a, and fig S5)**. Importantly, there was no significant response to TUG-1197 in FFA4-KO(*β*gal) mice (**fig 4a**).

**Fig 4.**
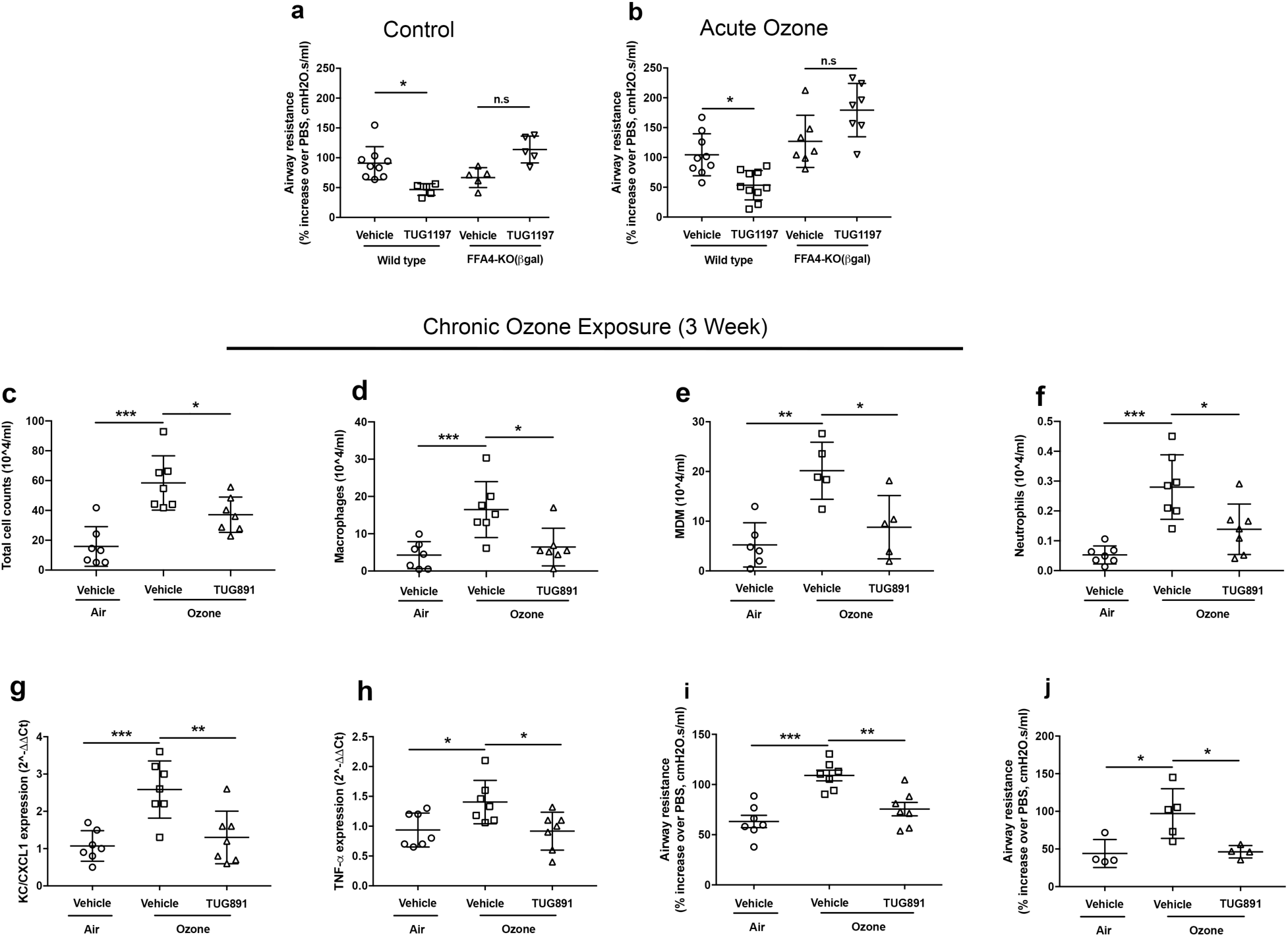
FFA4 agonism reduces airway resistance in healthy mice and produces broncho-relaxant, anti-inflammatory and prophylactic effects in ozone model of inflammatory lung disease. (a) Broncho-relaxant effect of TUG-1197 (0.364 mg/ml) on acetylcholine-induced (185 mg/ml) airway resistance in healthy wild type and FFA4-KO(βgal) mice. (b) Broncho-relaxant effect of TUG-1197 (0.364 mg/ml) on acetylcholine-induced (185 mg/ml) airway resistance in wild type and FFA4-KO(βgal) mice that had been exposed to ozone (3 ppm, 3 hr) 24 hours prior to the lung resistance experiments. (c-j) Anti-inflammatory effect of repeated administration of TUG-891 in a 3-week ozone exposure model. Mice were exposed with either control air or 3 ppm ozone for 3 hr, twice a week for 3 weeks. Before each exposure mice were treated with intranasal administration of either vehicle or TUG-891 (0.036 mg/ml). Cells in BAL fluid were obtained and enumerated for total cell counts (c) and analysed using flow cytometry to identify (d) alveolar macrophages, (e) monocyte derived macrophages (MDM) and (f) neutrophil populations. Lung tissue (left lobe) was isolated and RT-qPCR analysis was performed to identify changes in the expression of (g) KC/CXCL1 and (h) TNF-α genes. (i) Prophylactic and (j) broncho-relaxant effect of TUG-891 in 3-week ozone model where TUG-891 (0.036 mg/ml) was administered 1 hr prior to each ozone exposure (i) and at the end of the ozone exposure (j), during acetylcholine-induced (50 mg/ml) lung resistance measurements. Data presented are the mean ± S.E.M. of 4-7 animals. The data shown were analysed by Bonferroni multiple comparison test where *p<0.05, **p<0.01, ***p<0.001 and NS=not significant.

We next tested the effects of FFA4 activation within the context of an acute model of ozone pollution-induced inflammation. Under these conditions we observed no significant hyper-responsiveness induced by exposure to ozone but nevertheless administration of TUG-1197 was again seen to decrease airway resistance in wild type mice. This response to TUG-1197 was once more absent in FFA4-KO(*β*gal) mice (fig 4b). This was extended by testing the effects of FFA4 activation in a model of chronic (3 week) ozone exposure, which produced a profound inflammatory response as indicated by an increase in immune cells in broncho-alveolar lavage (BAL) fluid that included, alveolar macrophages, monocyte-derived macrophages and neutrophils, as well as up-regulation of the inflammatory mediators KC/CXCL1 and TNF*α* (**fig 4c-h**). This inflammatory response was accompanied by significant airway hyper-responsiveness (**fig 4i,j**). Under these conditions, administration of TUG-891, given 1 hr prior to each ozone exposure over the 3 week period, significantly decreased airway hyper-responsiveness (fig 4i) as well as decreasing airway inflammation as indicated by reduction of immune cells in the BAL fluid (**fig 4c-f**) and reduced levels of inflammatory markers (**fig 4 g-h**).

The effects of TUG-891 on neutrophil infiltration in the chronic ozone model prompted us to test the possibility that FFA4-induced anti-inflammatory effects might, at least in part, be mediated via modulation of leukotriene-mediated neutrophil infiltration. To assess this we analysed the transcript levels of 5-LOX, the enzyme responsible for LTB4 production, and the transcript levels of the BLT1 and BLT2 (BLT1 is the high affinity, and BLT2 the low affinity, LTB4 receptor). We found that mRNA encoding these genes was unchanged by 3 week ozone exposure and that TUG-891 treatment similarly had no effect on transcript levels **(fig S6a-c),** although there was a small trend towards a decrease of BLT1**. Interestingly, the levels of LTB4 itself were seen to increase with chronic ozone treatment and this increase was prevented by administration of TUG-891 (fig S6d).** That other factors may contribute to airway neutrophil infiltration, including increased production of the T helper 17 cell (TH17) interleukin IL17*α*^29–32^ was also considered, however, mRNA corresponding to this ligand was unchanged following chronic ozone exposure or treatment with TUG-891 **(fig S6e).**

Interestingly, FFA4 expression was detected in BAL cells that were also positively stained for each of CD11c, GR-1 and SiglecF **(fig S7).** These are markers respectively of monocytes, macrophages and dendritic cells, indicating that FFA4 activation might have a direct anti-inflammatory effect via these immune cells. It is also noteworthy that FFA4 was not detected in CD3^+^ or B220^+^ cells which are markers of T-lymphocytes which suggests that FFA4 is not expressed by T-lymphocytes **(fig S7).**

Whilst the above described data indicate that FFA4 has an anti-inflammatory role that could contribute indirectly to reduced hyper-responsiveness, results shown in **fig 3** on PCLS and in live mice in **fig 4a,b,** point to a more direct role for FFA4 on ASM relaxation. We therefore tested if acute FFA4 activation could reduce airway resistance in mice in which hyper-responsiveness had already been induced by chronic exposure to ozone. This potential was supported by data demonstrating that FFA4 transcript levels were not affected by chronic ozone exposure **(fig S8a)** nor was FFA4 signalling (calcium mobilization) altered **(fig S8b,c).** In settings in which hyper-responsiveness had been induced in mice by chronic ozone exposure, administration of TUG-891 together with acetylcholine significantly reduced airway resistance (**fig 4j**), indicating that FFA4 agonism can produce ASM relaxation, even in the context of an already established inflammatory lung disease.

In an alternative disease model we induced airway hyper-responsiveness by an 8-wk exposure to cigarette smoke (fig 5a). This procedure correlated with significant inflammation as indicated by an increase in numbers of aveolar macrophages and neutrophils (**fig 5b,c**). Acute administration of TUG-891 by nebulization prior to treatment with a bronchoconstrictor resulted in a reversal of hyper-responsiveness (**fig 5a**) indicating that, similar to the chronic ozone model, FFA4 activation in this cigarette smoke model was able to produce ASM relaxation.

**Fig 5.**
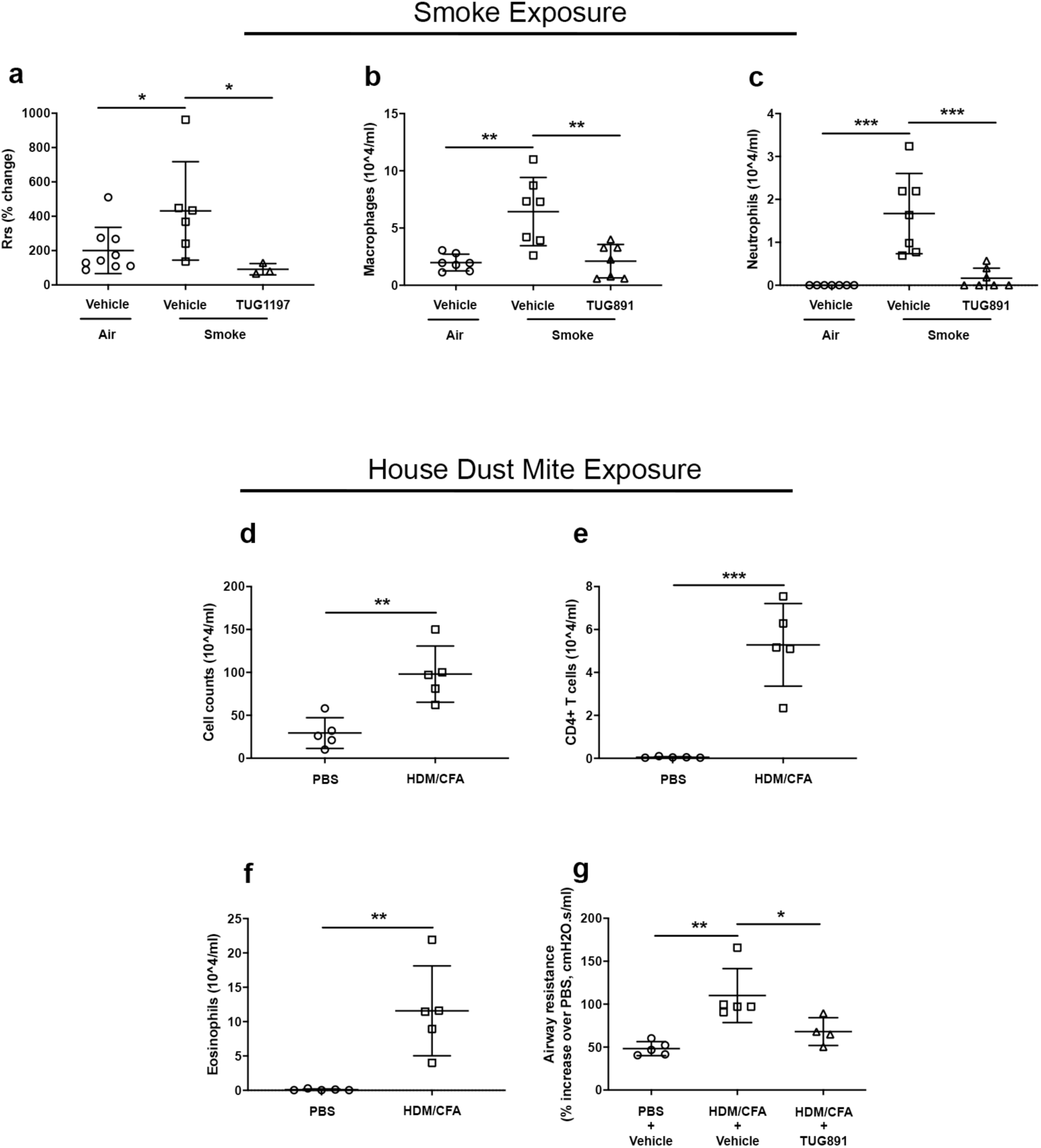
FFA4 agonism reduces increased airway resistance and lung inflammation induced by chronic cigarette exposure as well as airway resistance caused by exposure to house dust mites. Mice were exposed to air or cigarette smoke for 8 weeks. (a) Transpulmonary (Rrs) airway resistance was measured in air and smoke mice exposed to 3 mins of nebulized TUG-1197 (0.364 mg/ml) followed by 3 mins of nebulized methacholine (30 mg/ml). Mice were administered daily with TUG-891 (0.364 mg/ml) prior to each smoke exposure and the effect of chronic TUG891 dosing on (b) macrophages and **(c)** neutrophil infiltration (in BAL fluid) was measured. Mice were sensitised with subcutaneous injection of either PBS or house dust mite extracts supplemented with Freunds complete adjuvant (CFA, 1:1 mixture 100 μg HDM) and then challenged with intranasal delivery of PBS or HDM (25 μg). Mice were sacrificed and total cell counts **(d)** in BAL fluid were measured. CD4+ T cells (**e**) and eosinophils **(f)** were enumerated by flow cytometry. Mice were administered daily with either vehicle or TUG-891 (0.182 mg/ml) between the sensitization and final HDM challenge and the prophylactic effect **(g)** of TUG-891 on lung resistance was measured. Data represent the means ± S.D. of 3-9 animals where *p<0.05, **p<0.01 as determined by ANOVA with Bonferroni post-hoc test.

Importantly, FFA4 activation in the smoke model also appeared to be anti-inflammatory because chronic administration of TUG-891 prior to exposure to each cigarette also prevented airway inflammation as measured by a reduction in alveolar macrophages and neutrophils at termination of the study (**fig 5b,c**).

Finally, we induced airway hyper-responsiveness in mice by sensitization to house dust mites (HDM). Following sensitisation, immune cell count in the BAL fluid (**fig 5d**), and specifically CD4^+^ T cells (fig 5e) and eosinophils (**fig 5f**), was seen to increase, consistent with the notion that this model reproduces important features of allergic asthma^33^. Daily administration of TUG-891, after the initial sensitization and before the final HDM challenge, significantly reduced airway hyper-responsiveness (**fig 5g**). Alongside the ozone and cigarette-smoke models these data indicate that FFA4 activation can ameliorate airway inflammation and hyper-responsiveness in the context of multiple respiratory disease models.

### FFA4 is expressed in human airways and mediates bronchodilation

Transcript levels of FFA4 in human bronchial epithelial cells (HBEC) and human airway smooth muscle (ASM) derived from healthy donors was detected by PCR and qPCR (**fig 6a,b**). We confirmed the selectivity of a previously characterized, in-house human (h) selective FFA4 antiserum^21^ raised against a C-terminal peptide of hFFA4 (K^3^^42^-R^353^) in Western blots of lysates prepared from Chinese hamster ovary (CHO) cells transfected with hFFA4. In these experiments hFFA4 was detected as a broad band running at ∼50kDa only in lysates from hFFA4 transfected cells (**fig 6c**). This same antiserum detected hFFA4 in membrane and intracellular locations in immunofluorescent studies of hFFA4 transfected CHO cells (**fig 6d**). Using this antiserum we investigated the expression profile of hFFA4 in isolated HBEC and ASM by flow cytometry (**fig 6e,f**) and immunocytochemistry (**fig 6g,h**) as well as in immunohistochemistry performed on human bronchial biopsies (**fig 6i-l and fig S9a,b**). These experiments established that hFFA4 was primarily expressed in the human airway epithelium with lower levels in the ASM.

**Fig 6.**
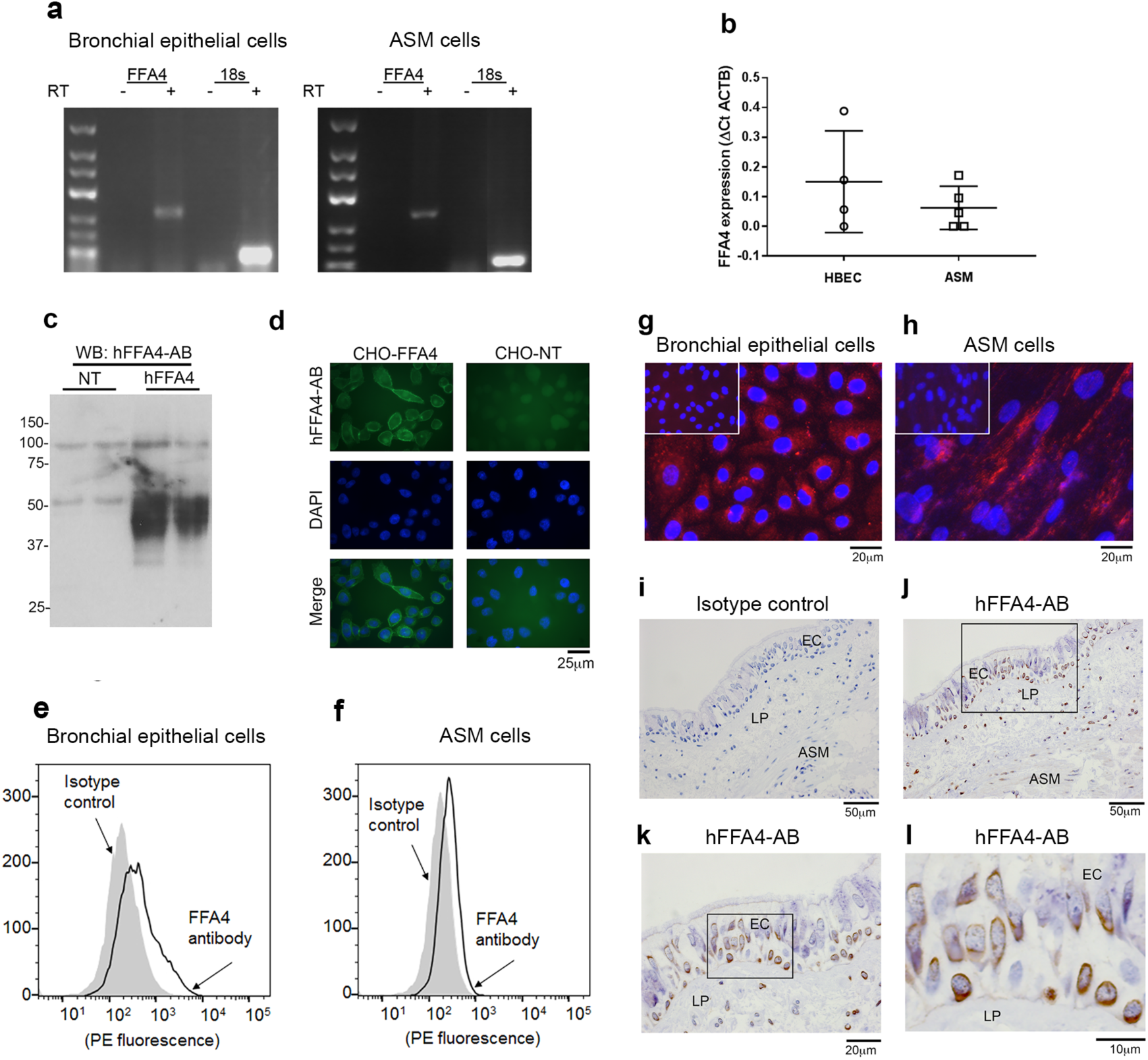
FFA4 is expressed in human bronchial epithelial cells and human smooth muscle cells and can be detected at both transcript and protein levels. **(a)** Transcript levels of FFA4 and 18s (as a control) were determined in bronchial epithelial cells (n=5 donors) and ASM cells (n=5 donors) using RT-PCR. Shown is a representative gel. The expected sizes of PCR products are as follow; FFA4 251bp,18s 93 bp. **(b)** Quantitative RT-PCR of bronchial epithelial cells (4 donors) and ASM cells (5 donors) was conducted using ACTB housekeeping gene as a control. Data represent mean ± SEM. **(c)** Western blot of human FFA4 transiently transfected in HEK293 cells using an antiserum selective for FFA4 developed in-house (hFFA4-AB). **(d)** Immunofluorescence images of human FFA4 stably transfected in CHO-Flip-In cells (CHO-FFA4) and control non-transfected CHO cells (CHO-NT). **(e-f)** Example fluorescent histogram of FFA4 expression (black trace) in bronchial epithelial cells and ASM by flow cytometry verses isotype control antibody (gray shading); fold increase in geometric mean fluorescence intensity (GMFI) of anti-FFA4/isotype control antibody (95% CI) 3.019 (2.042-3.996) [11 donors, P<0.01] and ASM cells 2.241 (1.662-2.819) [14 donors, P<0.001] two tailed t test against isotype control. **(g,h)** Representative photomicrograph (x40 magnification) showing bronchial epithelial cells (5 donors) FFA4 expression (red, left; isotype control antibody insert) and ASM cell (4 donors) FFA4 expression (red, right; isotype control insert) nuclei stained blue with 4’,6-diamidino-2-phenylindino-2-phenylindole blue by immunofluorescence. **(i-l)** Representative photomicrographs of normal human bronchial biopsy sections stained with **(i)** isotype control antiserum, **(j)** human FFA4 selective antiserum (4 µg/ml) and **(k)** zoom of selected region of the epithelium from **(l)** EC = epithelial cell layer, ASM = airway smooth muscle. LP = Lamina propria.

The expression profile observed in human lung closely mirrored that seen in the mouse lung (see; **fig 1c,d**), further suggesting that the mouse airway responses reported here might mimic those of hFFA4 in human airways. This possibility was tested further by monitoring calcium mobilization in response to TUG-891 in human ASM and HBECs. TUG-891 (50 μM) caused a statistically significant increase in calcium mobilization in both ASM and HBEC cells (**fig 7a,b**). We next assessed if FFA4 in human ASM might mediate relaxation using a collagen gel preparation where human ASM contraction can be monitored by reduction in the diameter of the collagen gel^34^. Treatment of ASM in this preparation with TUG-891 reversed both spontaneous contraction and contraction observed in the presence of the bronchoconstrictor, acetylcholine (**fig 7c,d**). These data demonstrate that FFA4 is expressed and functional in isolated human ASM cells and that activation of this receptor is able to promote human ASM relaxation *ex vivo*.

**Fig 7:**
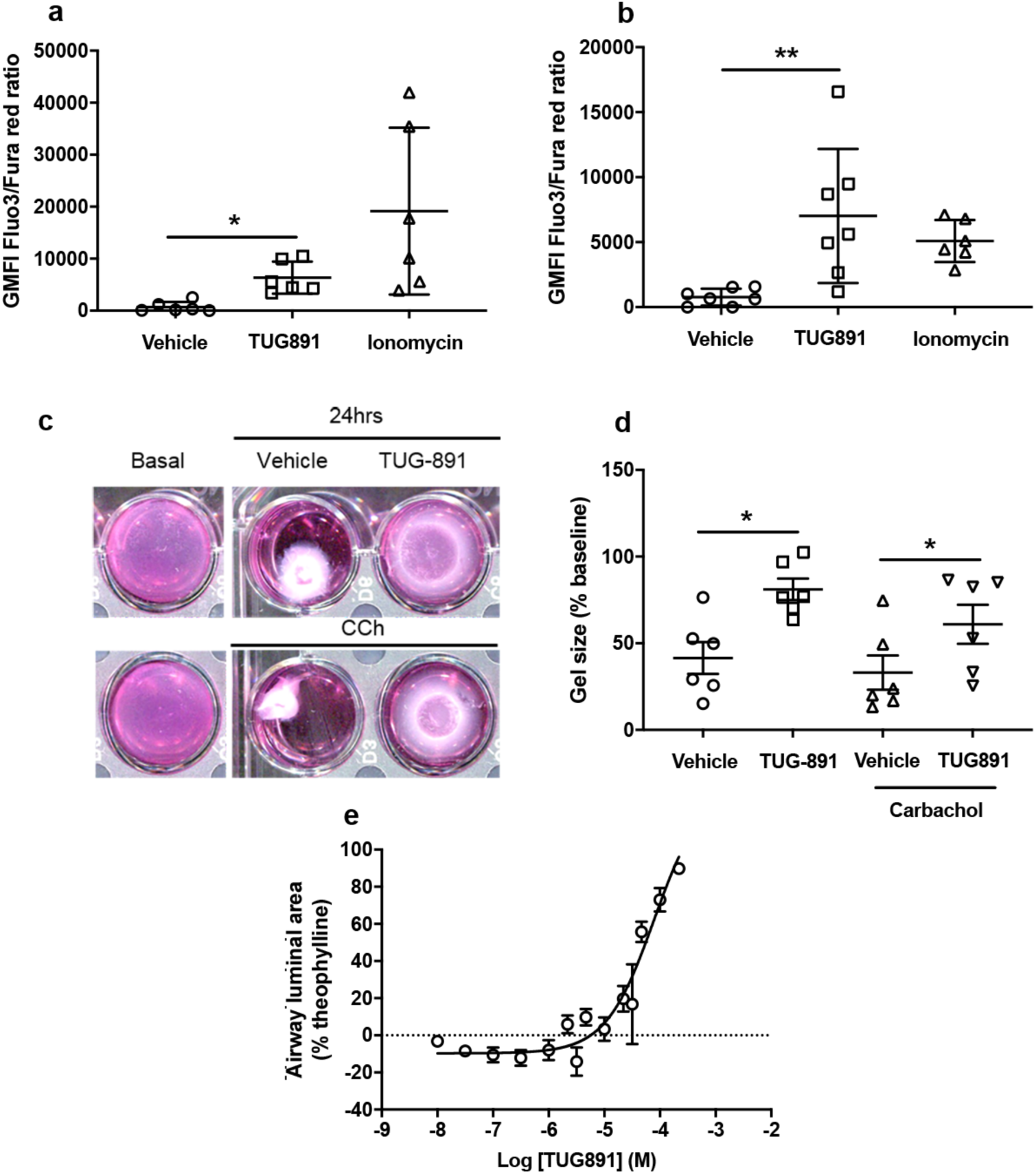
Functional responses of FFA4 in isolated human lung tissues and cells. **(a)** Intracellular calcium (iCa^2+^) elevation in bronchial epithelial cells (6 donors) and **(b)** ASM cells (6 donors) in response to TUG-891 treatment or ionomycin (1.5μg/ml) as a positive control. GMFI equates to total stimulated GMFI minus matched baseline minus vehicle control. Data are plotted as mean ±SEM. **(c)** Representative gel photographs taken at 0 hours (basal) and 24 hours either uncontracted or pre contracted with carbachol (CCh 100 μM) followed by treatment of TUG-891 (50μM). **(d)** Percentage contraction of collagen gels in the presence of ASM cells (6 donors) either uncontracted or pre contracted with carbachol (CCh 100 μM) followed by treatment of TUG-891 (50 μM). Data are plotted as mean ± S.E.M. **(e)** Concentration-response data for TUG-891 mediated relaxation of human airway strips pre-contracted with carbachol (100 μM). The data presented in **e** are the mean ± S.E.M of at least three independent experiments from healthy donors.

We extended these studies by testing human ASM contractile responses in intact lung tissue (un-denuded bronchi) isolated from healthy subjects using wire myography. In these experiments TUG-891 produced a concentration-dependent (pEC_50_, 4.34 ± 0.13) relaxation of human airways that had been pre-contracted with acetylcholine (**fig 7e**).

### Potential role of prostaglandins in the mechanism of FFA4-mediated bronchodilation

It has been known for some time that through an interplay between EP_1_ (contraction) and EP_2_ (relaxation) prostanoid receptors, prostaglandins can regulate lung function, via both contraction and relaxation of ASM^35^. Among the prostanoids reported to have an impact on lung function is prostaglandin PGE_2_ ^35^. Released from various lung cell types including ASM ^36–38^, airway epithelium ^39^ and immune/inflammatory cells of the lung ^40^, PGE_2_ can mediate bronchodilation via EP_2_ receptors^41–43^. Interestingly, the bronchodilation properties of a number of GPCR ligands including bradykinin, substance P, ATP, and proteases that activate the proteinase activated receptors (i.e. PAR2), are mediated by the release of PGE_2_ ^37, 44, 45^. Therefore, in addition to directly acting on ASM we tested the possibility that FFA4 might also mediate bronchodilation via the release of PGE_2_. Incubation of PCLS derived from wild type mice with TUG-891 resulted in a significant release of PGE_2_ whereas the same concentration of TUG-891 applied to PCLS from FFA4-KO(*β*gal) mice generated no significant PGE_2_ release (**fig 8a**). Importantly, we confirmed that PGE_2_ administration to pre-contracted PCLS derived from wild type mice resulted in relaxation of ASM (**fig 8b**). Moreover, the EP_2_ selective antagonist, PF-04418948 ^46^, significantly blocked the relaxation of ASM in response to TUG-891 in PCLS (**fig 8c**). TUG-891 was used in this set of experiments since our *in vitro* studies indicated that it is the more potent of the two compounds (**fig 2a,c,e**). Finally we measured PGE_2_ levels in the BALF from mice exposed to ozone and treated with TUG891. In these experiments we observed a significant increase in PGE_2_ following chronic administration of TUG891 (**fig 8d**).

**Fig 8:**
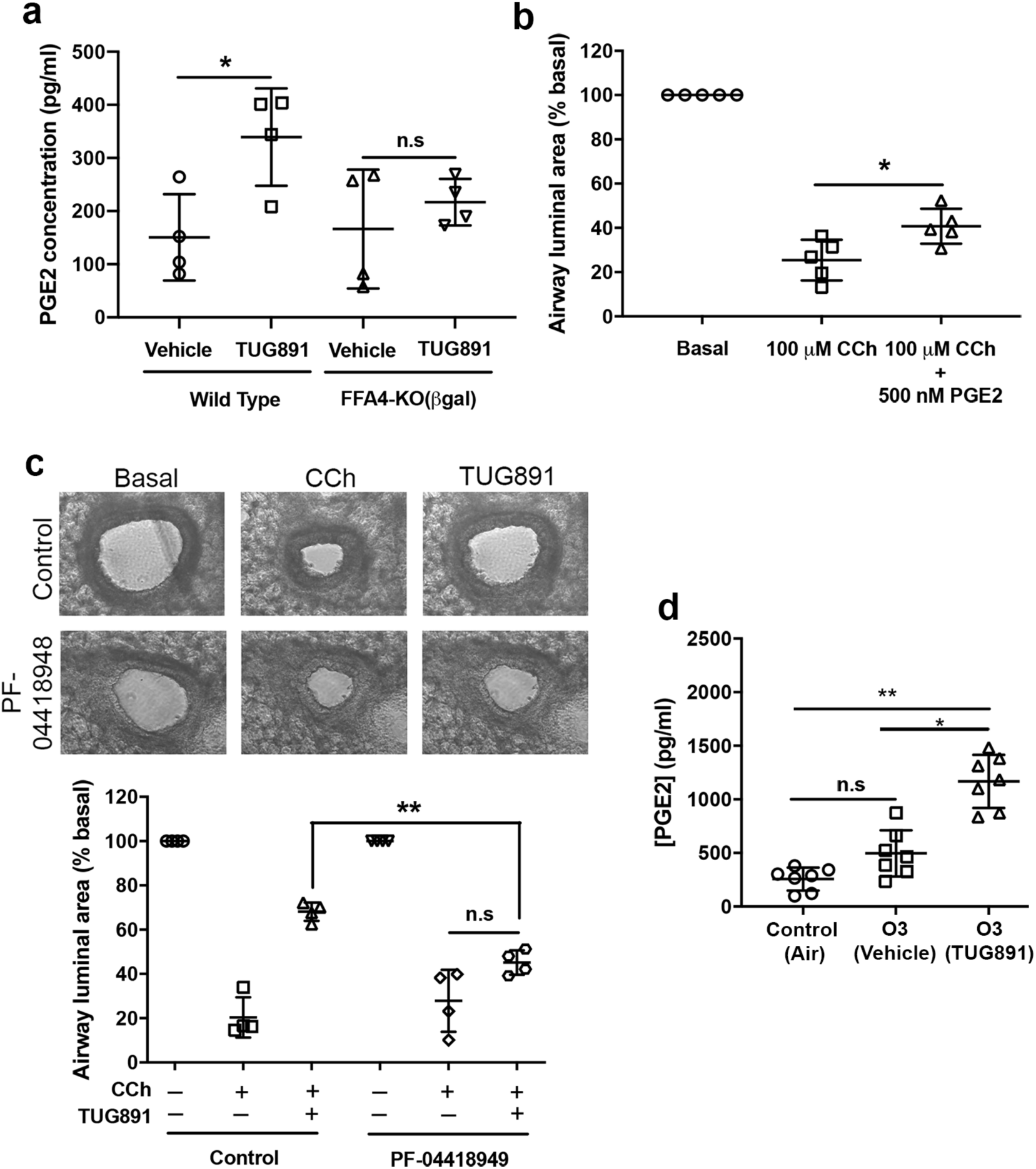
FFA4 mediated airway relaxation in mouse is partly dependent on PGE_2_ and the prostanoid receptor EP_2_. **(a)** PGE_2_ release from PCLS obtained from the lungs of wild-type mice and FFA4 KO(βgal) mice was detected by ELISA. **(b)** Effects of PGE_2_ (500 nM) treatment of carbachol (CCh) pre-contracted PCLS. **(c)** Effect of pre-treatment of PCLS with the selective EP_2_ receptor antagonist PF-04418948 (10 μM) for 1 hr on TUG-891 response. **(d)** PGE_2_ release into the bronchoalveolar lavage fluids (BALFs) of mice exposed to normal air or chronically exposed to ozone treated with either vehicle (1% DMSO) or TUG891 (100 μM). Data represent the means ± S.D. of at least 3 animals where *p<0.05, **p<0.01, NS = not significant as determined by ANOVA with Bonferroni post-hoc test.

Previous work has highlighted species differences in the roles of prostanoid receptor subtypes in mediating PGE_2_-dependent airway relaxation, with EP_2_ being the predominant subtype in the mouse and EP_4_ the key subtype in human^47, 48^. We confirmed here expression of all four known EP prostanoid receptors (EP_1-4_) in isolated human bronchial epithelial cells **(fig S10a).** However, only EP_2-4_ were expressed in human ASM **(fig S10b).**

To further investigate the mechanism by which FFA4 mediates increased PGE_2_ production in mouse, gene expression analysis was performed in lung tissues obtained from the 3-wk ozone exposure model. In this experiment, mRNA levels of the majority of genes involved in the biosynthesis of PGE_2_, including PGE_2_ synthases and cyclooxygenase-1 (COX1) were not significantly altered whilst there was a small, but significant, up-regulation of cyclooxygenase-2 (COX2) mRNA **(fig S11 a-e).** This suggests that FFA4 activation may induce expression of COX2 to increase PGE_2_ levels. However, further work would be required to assess the significance of this observation not least because other mechanisms might be in play to regulate PGE_2_ levels such as availability of arachidonate and post-translational regulation of synthetic enzyme activity. Importantly, in the 3-wk ozone exposure model expression of the EP_2_ receptor was not significantly altered **(fig S11 f),** highlighting that this pathway would still be expected to be operational in the disease state.

Because the FFA4 agonists appeared to produce a response via EP_2_ receptors we assessed if either TUG-891 or TUG-1197, which represent very different chemotypes, could directly activate EP_2_ receptors. In these experiments TUG-891 (10 μM) produced no significant activation of the EP_2_ receptor (**table S1**). TUG-891 also lacked agonist function at other EP prostanoid receptors, including the EP_1_ receptor (**table S1**). Equally TUG-891 did not significantly block effects of endogenously generated prostanoids at any of the EP prostanoid receptors (**table S2**). A similar lack of effect of 10 μM TUG-1197 as an agonist (**table S1**) or antagonist (**table S2**) on each of the EP_1-4_ prostanoid receptors was recorded. These data provide further support that the effect of ligands with potency at FFA4, in both *ex vivo* and *in vivo* studies, is indeed mediated by FFA4 (and not via off-target activity at, for example, prostanoid receptors).

Since we established a link here between FFA4 bronchodilation and PGE_2_ we wanted to test if PGE_2_ might also be involved in the FFA4-mediated anti-inflammatory response observed in mouse disease models. That this might be the case was indicated by the observation that PGE_2_ levels were increased in BALF following chronic administration of a FFA4 agonist in the ozone model (fig 8d) - a response that correlated with an increase in COX2 transcription (fig S11 a-e), the enzyme response for prostaglandin biosynthesis including PGE_2_. In an attempt to investigate this possibility we tested the effects of the COX2 inhibitor celecoxib on the FFA4-mediated anti-inflammatory response by monitoring changes in number of cells, TNF*α* levels and IL-1*β* transcript in BALF from mice chronically exposed to ozone. In these experiments celecoxib alone was anti-inflammatory which phenocopied TUG-891 anti-inflammatory response (fig S12 a-c). The co-administration of celecoxib and TUG-891 provided an anti-inflammation effect equivalent to celecoxib and TUG-891 alone (fig S12 a-c). Hence, unlike the bronchodilation response where it was possible to define a role for PGE_2_ in the FFA4 response we were not able to conclude from these experiments a specific role for PGE_2_ in FFA4-mediated anti-inflammation.

## Discussion

We present evidence that the GPCR, FFA4, is expressed and active in the murine lung, where activation of the receptor mediates ASM relaxation under both normal physiological conditions and in the context of respiratory disease models whilst, additionally, generating an anti-inflammatory effect. That these mouse studies have applicability to human disease is supported by data establishing FFA4 expression and function in human ASM and bronchial epithelial cells (HBEC) and in producing relaxation of human ASM in response to the agonist TUG-891. Together, our data support the conclusion that the long chain free fatty acid receptor FFA4 might be a tractable pharmacological target for the treatment of human airway diseases.

It is important to note the FFA4 selective agonist TUG-1197, and the FFA1/FFA4 co-agonist TUG-891, were equally effective in mouse lung *ex vivo* and *in vivo* experiments. However, in human cell and tissue-based studies we found that TUG-891 was more effective. Although this might be explained by the fact that TUG-891 is slightly more potent than TUG-1197 at FFA4 it might also suggest that, in addition to FFA4, FFA1 may also contribute to the airway responses reported here, particularly in human lung where the expression of functional FFA1 has been reported ^49^. In contrast to our data however an ex vivo study has suggested that FFA1 might augment acetylcholine mediated ASM contraction rather than mediate ASM relaxation ^49^. Since our data supports the opposite possibility we would argue that there is a potential for dual FFA1/FFA4 agonists to be more effective agents at mediating ASM relaxation than FFA4 selective agonists. Hence, testing dual agonists alongside FFA4 selective agents in experimental medicine studies will determine which approach is desirable ahead of human clinical trials. It will also be important to reduce the possibility of off-target adverse responses before human trials by significantly increasing the potency of our current tool compounds. That off-target activity of our current ligands may exist is indicated by the small calcium transient (see; **fig 2i**) and trend (although not significant) relaxation response to TUG-1197 in PCLS derived from FFA4-KO(*β*gal) mice (see; **fig 3h**).

We provide evidence that one mechanism of action of FFA4 in promoting ASM relaxation in mouse is via receptor-mediated release of the prostaglandin PGE_2_ that subsequently acts at the EP_2_ prostanoid receptor to mediate relaxation. A similar mechanism of ASM relaxation has been reported for other GPCRs ^37, 44, 45^ raising the question of whether an alternative approach in a therapeutic context might be to directly target EP_2_ to deliver ASM relaxation. However, prostanoid biology in the lung is complex. For example, EP_4_ is considered as the most predominant subtype in human^47, 50^ and therefore targeting specifically EP_2_ would be less likely to be effective in human. Moreover, PGE_2_ has multiple modes of action in that it can activate four different receptor subtypes (EP_1_-EP_4_) which transduce a variety of processes including bronchodilation, cough, microvasculature leaks and regulation of pulmonary blood vessel tone^48, 51, 52^. Our data also show that mRNA encoding each of the four subtypes can be detected in human ASM and/or bronchial epithelial cells. As such, the indirect generation and action of PGE_2_ produced via activation of FFA4 may contribute to the highly beneficial outcomes observed via a number of receptors, whilst a selective agonist for one or other subtype might fail to produce the integration of outcomes generated by the native prostanoid.

Our study points to FFA4 having a dual effect in the context of inflammatory airway disease. The first centres on an action on the ASM resulting in bronchodilation with the potential of symptomatic relief in conditions such as asthma where airway constriction is a feature. Although our studies on human airways (collagen gel contraction) point to a direct effect of FFA4 on ASM the relatively high levels of expression of this receptor on epithelial cells means that an indirect effect cannot be ruled out. It is possible for example that the airway epithelium is the origin of FFA4-mediated PGE_2_ release that subsequently leads to ASM relaxation. Secondly, our data point to a role for FFA4 in the regulation of inflammation that is consistent with reports that FFA4, expressed on various immune cells including macrophages ^14, 53^, can act as a negative modulator of inflammation in the context of metabolic disease ^14, 15, 54^. Exposure of mice to repeated doses of ozone resulted in a profound inflammatory response, evidenced by an increase in immune cells (neutrophils and macrophages) in the airway spaces and a corresponding elevation of gene expression for the macrophage pro-inflammatory cytokine TNFα, and the neutrophil chemotactic factor, KC/CXCL1. Importantly, intranasal pre-treatment of mice with TUG-891 prior to each exposure to ozone significantly reduced the expression of these pro-inflammatory cytokines and prevented immune cell infiltration. Thus, our data indicates that in addition to an effect on ASM contraction that has the potential for symptomatic treatment of inflammatory airway disease, FFA4 activation could also have an anti-inflammatory property that would provide for disease modification.

It is not clear from our study whether the dual action of FFA4, namely relaxation of ASM and anti-inflammation, is via a shared mechanism. For example, we provide evidence that ASM relaxation is in part mediated by the release of PGE_2_ acting on EP_2_ receptors however we currently have no evidence for the involvement PGE_2_ release in the anti-inflammatory response. The detection of FFA4 expression in CD11c, GR-1 and SiglecF positive immune cells in the lung might indicate that FFA4 has a direct role in regulating immune cell response and therefore a distinct anti-inflammatory mechanism from that which mediates ASM relaxation. Despite these unresolved questions regarding the mechanism and cell types involved in mediating FFA4 dual effects on ASM and inflammation our study reveals the therapeutic potential of targeting FFA4 in inflammatory lung disease.

A further important question that remains to be addressed is the origin of endogenous LCFAs that regulate lung FFA4. It is noteworthy that we observed no significant difference in the ‘basal’ airway resistance between wild type and FFA4-KO mice (see; fig 4a) indicating that more sensitive approaches and possibly additional mouse models that modulate endogenous lung fatty acid metabolism will be required to fully appreciate the role of endogenous free fatty acids. However, the omega-3 fatty acids, eicosapentaenoic acid (EPA) and docosahexaenoic acid (DHA) have been reported to be enriched in airway mucosa^55^ where they are proposed to act as precursors for the biogenesis of lipid mediators such as the resolvins that resolve inflammatory responses^56^, including lung inflammation^30^ via interacting with members of the GPCR family including the chemerin-1 and chemerin-2 receptors. Whether this enriched pool of dietary-derived omega-3 fatty acids, that can potentially be augmented by the recruitment of circulating DHA and EPA at the sites of inflammation^57^ or that can be released from membranes by the activity of phospholipase A_2_^58^, might be the source of endogenous LCFA activators for FFA4 is a possibility that is currently under investigation.

Importantly, our data demonstrated that expression of FFA4 in human airways is similar to that observed in mouse and that FFA4 agonism promotes relaxation isolated human ASM and of human airway strips. In this way we provide unique proof-of-concept that pharmacological activation of lung FFA4 can generate *in vivo* efficacy. The precise pharmacological properties of a clinically effective hFFA4 agonist, particularly whether a full/partial agonist or whether a dual FFA4/FFA1 agonist would be most efficacious or indeed whether a G protein-biased ligand that might avoid potential challenges such as receptor desensitization and tachyphalaxis^20^, is preferable has yet to be fully established. Our study does however provide the first evidence that FFA4 can indeed promote airway relaxation and resolve airway inflammation, and thus may be a *“druggable”* target for the treatment of airway diseases associated with bronchoconstriction and inflammation.

## Materials and Methods Ozone exposure

For the acute ozone exposure protocol, wild-type adult male C57BL6/N mice (8-12 wk old) FFA4-KO(*β*gal) mice (a kind gift from AstraZeneca as described previously ^59^ were administered with either medical air or 3 parts per million (ppm) ozone for 3 hr in a sealed Perspex container (EMB104, EMMS). Ozone concentration level of 3 ppm was continuously monitored with an ozone probe (ATi Technologies, Ashton-U-Lyne, UK). 24 hr after exposure, mice were used for experiments. For sub-chronic treatment, mice were administered intranasally with either vehicle or FFA4 agonist (TUG-891, 100 μM, 25 μl volume) 1 hr prior to ozone or control air treatments. Mice received 3 parts per million (ppm) ozone produced from an ozoniser (Model 500 Sander Ozoniser, Germany) for 3 hours, twice a week, for a total duration of 3 weeks. Control animals received filtered air only for the equivalent exposure period. In some experiments, mice were exposed to ozone for 3 weeks and then administered TUG-891 at the end, during the lung function measurements.

## House dust mite mouse model

Adult male Balb/c mice (8-12 wk old from Envigo, Oxford, UK) were sensitized on day 0 with subcutaneous injection of 100 μg in 100 μl volume of house dust mite extract (HDM, Citeq Biologics, Groningen, The Netherlands) supplemented with Freund’s Complete Adjuvant (1:1 mixture, Sigma, Dorset, UK). Control mice were given an injection with equivalent volume of PBS (Life Technologies, Paisley, UK). Mice were then administered TUG-891 daily (500 μM, 25 μl volume) for 15 days via intranasal delivery under general anaesthetic. On day 15 mice were challenged with 25 μg of HDM in 25 μl volume delivered intranasally under anaesthetic condition. Mice were used for experiments 48 hrs post challenge.

## Supplementary Material

### Materials and Methods

#### Cigarette mouse model

6-8-week-old female wild type mice (Australian BioResources, NSW) were exposed to twelve 3R4F cigarettes (University of Kentucky) using a custom-designed and purpose-build nose-only smoke system (CH Technologies) twice per day, 5 times per week for 8 weeks, or room air as previously described ^60–62^. In some experiments, 100 μM TUG-891 was administered daily prior to cigarette smoke exposure.

#### Assessment of cigarette mouse model lung function

Mice were anesthetized (50 μl/10 g i.p.) with a mixture of xylazine (2 mg/ml, Troy Laboratories) and ketamine (40 mg/ml, Ceva). Mice were cannulated *via* the trachea and ventilated with a tidal volume of 8 ml/kg at a rate of 450 breaths/min. FlexiVent^TM^ apparatus (FX1 System; SCIREQ, Montreal, Canada) was used to assess airway resistance as previously described ^63–65^. Mice were ventilated for 5 mins before baseline recordings were obtained using single frequency (2.5Hz, 1.2s long; Snapshot-150) and broadband low frequency (1-20.5 Hz, 8s long; Quick-Prime-8) forced oscillation manoeuvres. Mice were then challenged with aerosolised PBS for 1 min and Snapshot-150 and Quick-Prime-8 recorded, followed by aerosolised TUG-1197 (0.364 mg/ml) for 3 mins (Snapshot-150 and Quick-Prime-8 recorded), and then aerosolised methacholine (30 mg/ml, Sigma) for 3 mins (Snapshot-150 and Quick-Prime-8 recorded). Rrs (from Snapshot-150) and Rn (from Quick-Prime-8) were analysed as a percentage change from TUG-1197.

#### Bronchoalveolar lavage

Bronchoalveolar lavage (BAL) fluids were obtained by introduction of two times 800 µL PBS via a cannula introduced into the trachea. BAL fluids so recovered were centrifuged at 500g for 10 min at 4°C. Cell pellets were resuspended in PBS supplemented with 1% FBS and 1 mM EDTA, mixed with 4% trypan blue solution and counted on a Countess (Life Technologies, Pisley, UK) automated cell counter for total cell numbers. Remaining cells were used for flow cytometry.

#### Immunohistochemistry of human lung biopsies

Bronchial biopsy specimens from healthy subjects were processed and embedded as described in^66^. 2 micron sections were then processed on the Autostainer Link 48 platform (Dako AS480) using an automated staining protocol for the FFA4 antiserum (4 µg/ml) and its rabbit polyclonal isotype control (Immunostep, Salamanca, Spain) using EnVision FLEX kit (Dako). Pictures were obtained using a Carl Zeiss Imager Z2 microscope and AxioCam HRc digital camera (Carl Zeiss, Germany).

#### Flow cytometry

Cells obtained from BAL were resuspended in FACS buffer (PBS supplemented with 1% FBS and 1 mM EDTA) and then stained with CD4, CD11b, CD11c, CD64, Siglec-F, MerTK, F4/80 and Ly6G antibodies (obtained from either Biolegend, London, UK or Life Technology, Paisley, UK) for 30 min at 4°C. Cells were washed 3x with 1 ml FACS buffer and finally resuspended in 300 μL of FACS buffer for immunophenotyping. Staining for FFA4 expression in mouse BAL cells or human bronchial epithelial cells and ASM cells were performed in 0.5% triton-x100 permeabilised cells using in house anti-FFA4 antisera followed by fluorescently labelled secondary antibody (Bio-Rad, Watford, UK). Rabbit isotype control (Immunostep, Salamanca, Spain) was used for definition of non-specific staining. BAL samples that were contaminated with red blood cells were treated with e-bioscience rbc lysis buffer (Life Technologies, Paisley, UK) prior to staining. Alveolar macrophages were identified as Siglec-F+, CD11c+, CD11b^int^ and F4/80+ whereas gatings for neutrophils were Siglec-F-, CD11c-, CD11b+ as Ly6G+. Monocyte-derived macrophages were classed as CD64^high^, CD11c^high^, F4/80+, MerTK^+^, SiglecF^low^. Eosinophils were identified as being SiglecF+, CD11c- and CD4 positive whilst T cells were identified based on positive staining for CD4 glycoprotein. Flow cytometry was performed with LSRII or FACSCanto (BD Biosciences) instruments and data were analysed with FlowJo vX software (FlowJo LLC).

#### BRET arrestin3 recruitment assay

To assess arrestin3 recruitment to the FFA4 receptor, a bioluminescence resonance energy transfer (BRET) assay was used based on protocols previously described ^3, 18, 21^. Briefly, HEK293T cells were co-transfected with arrestin3-Rluc and FLAG-FFA4-eYFP plasmids in a 1:4 ratio using polyethyleneimine. After 24 hr incubation, cells were sub-cultured into poly-d-lysine-coated white 96-well microplates and incubated for a further 24 hr prior to the assay. Cells were then washed with Hanks’ balanced salt solution (HBSS) and incubated in HBSS for 30 min prior to conducting the assay. To initiate the assay, the Rluc substrate coelenterazine-h was added to a final concentration of 2.5 μM and incubated for 10 min at 37 °C before the test compound was added. Following a further 5-min incubation, luminescence emissions at 535 and 475 nm were measured using a PHERAstar FS (BMG Labtech, Offenburg, Germany), and the BRET signal was represented as the 535/475 ratio multiplied by 1000 to yield the arbitrary milli-BRET units (mBRET).

#### Ca^2+^ signalling in cell lines, ASM and HBEC

Ca^2+^ assays were carried out on Flp-In T-REx 293 cells that inducibly express either FFA4 or FFA1 receptor upon treatment with doxycycline. Cells were seeded at 50 000 cells/well in black clear-bottom 96-well plates and allowed to adhere for 3-4 hr in a humidified 37°C incubator at 5% CO_2_, 95% air. Cells were then treated with 100 ng/ml doxycycline overnight to induce receptor expression. The following day, cells were incubated in culture medium containing the Ca^2+^-sensitive dye, FURA2-AM (3 μM) for 45 min. Cells were then washed three times to remove fatty acids present in the culture medium and then allowed to equilibrate for 15 min in HBSS before conducting the assay. FURA2 fluorescent emission was measured at 510 nm following excitation at both 340 and 380nm using a Flexstation plate reader (Molecular Devices). Ca^2+^ responses were measured as the difference between 340:380 ratios before and after the addition of appropriate ligands. Alternatively, ASM and HBEC incubated in 500 μl of pre-warmed PBS (supplemented with 2mM Ca2+, pH7.4) containing 10 μg/ml fura-red (Invitrogen, Paisley, UK) and 4 μg/ml fluo-3 green (Invitrogen, Paisley, UK) at 37°C, 5% CO_2_ for 45 minutes in the dark. A further 500μl of pre-warmed PSS was added to the cells prior to centrifugation at 1300rpm for 8 minutes at 20°C. Supernatants were discarded and cells were resuspended in 300μl PBS in FACS tube. Samples were analysed using a BD FACSCanto flow cytometer. Baseline recordings were performed for 1 minute. Following this vehicle control, TUG-891 (30-100 µM) or the positive control; ionomycin (1.5 μg/ml, Sigma), was added to the cells and recording continued for a further 3 minutes. Ratiometric analysis of the geometric mean fluorescence of fluo-3 green/fura-red was conducted as an indication of mean relative intracellular Ca^2+^ (Ca ^2+^i) concentration over a period of 3 minutes post-stimulation. Data are plotted as mean± SEM.

#### Reverse Transcriptase PCR

Wild type female C57BL/6N or FFA4-KO(*β*gal) adult mice (8-12 wk old) were sacrificed by cervical dislocation followed by exsanguination. Lung tissue was isolated and total RNA was prepared from 100 mg of the tissue using an RNeasy Lipid Tissue Mini Kit (Qiagen) as per the manufacturer’s instructions. For human bronchial epithelial cells and ASM, total RNA was isolated using the peqGOLD total RNA kit (Peqlab, VWR). RNA concentration was verified using the nanodrop 2000 spectrophotometer (ThermoFisher, Waltham, USA). Complementary DNA synthesis was performed using SuperScript III First-Strand Synthesis SuperMix (Life Technologies) with 1 μg of total RNA template per reaction. Phusion DNA polymerase (New England Biolabs) was used in PCR reactions to amplify the gene for mouse FFA1 receptor, mouse FFA4 receptor and α-tubulin using 2 μL of first-strand synthesis reaction products as a template. The following PCR program was used: 95°C 3 minutes, 95°C 30 seconds, 60°C 30 seconds, 72°C 30 seconds (40 cycles of three last steps). PCR products were then analysed by electrophoresis on 2% agarose gel. The following primer pairs were used:

Mouse FFA1 Forward: 5’-ACCTGCCCCCACAGTTCTCCTT-3’ Mouse FFA1 Reverse: 5’-AGAGTCGGGATCCCAGGCTT -3’ Mouse FFA4 Forward: 5’-ATGTCCCCTGAGTGTGCACA-3’ Mouse FFA4 Reverse: 5’-AAGAGCGTGCGGAAGAGTCG-3’

Mouse α-tubulin Forward: 5’-ATGCGTGAGTGCATCTCCATCCAT-3’

Mouse α-tubulin Reverse: 5’-AACATTCAGGGCCCCATCAA-3’ Human FFA4 forward: 5’-CCTGGAGAGATCTCGTGGGA -3’

Human FFA4 reverse: 5’- AGGAGGTGTTCCGAGTCTGG -3’

Human EP1_forward: 5’- GGTATCATGGTGGTGTCGTG -3’

Human EP1_reverse: 5’- GGCCTCTGGTTGTGCTTAGA -3’

Human EP2_forward: 5’- CCACCTCATTCTCCTGGCTA -3’

Human EP2_reverse: 5’- CGACAACAGAGGACTGAACG -3’

Human EP3_forward: 5’- CTTCGCATAACTGGGGCAAC -3’

Human EP3_reverse: 5’- TCTCCGTGTGTGTCTTGCAG -3’

Human EP4_forward: 5’- TGGTATGTGGGCTGGCTG -3’

Human EP4_reverse: 5’- GAGGACGGTGGCGAGAAT -3’

#### Quantitative RT-PCR

Wild type C57BL/6N or FFA4-KO(*β*gal) adult female mice (8-10 wk old) were sacrificed by cervical dislocation followed by exsanguination. Lung tissue was isolated and total RNA was prepared from 100 mg of the tissue using an RNeasy Mini kit (Qiagen) following the manufacturer’s instructions. cDNA was subsequently obtained using up to 1 µg of RNA using the QuantiTect Reverse Transcription Kit (Qiagen). For human bronchial epithelial cells and ASM, total RNA was isolated using the peqGOLD total RNA kit (Peqlab, VWR). RNA concentration was verified using the nanodrop 2000 spectrophotometer (ThermoFisher, Waltham, USA). 2 μL of the resulting cDNA were used in qPCR reactions set up with the following probes: mouse FFA4 (Mm00725193_m1), mouse FFA1 (Mm00809442_s1), mouse GAPDH (Mm99999915_g1), human FFA4 (Hs00699184_m1) and human ACTB (Hs99999903_m1) (Life Technologies) using the 7900HT Fast Real-Time PCR System (Applied Biosystems).SYBR green-based reactions were also carried out using the following primers: Mm_Ptges_1_SG QuantiTect Primer Assay (QT00118223), Mm_Ptges2_1_SG QuantiTect Primer Assay (QT00495348), Mm_Ptges3_1_SG QuantiTect Primer Assay (QT01759387), Mm_Il17a_1_SG QuantiTect Primer Assay (QT00103278), mHPRT forward

5’-AGGCCAGACTTTGTTGGATTTGAA-3’, mHPRT reverse

5’-CAACTTGCGCTCATCTTAGGCTTT-3’, mKC/CXCL1 forward

5’- CGCTGCTGCTGCTGGCCACCA-3’, mKC/CXCL1 reverse

5’-GGCTATGACTTCGGTTTGGGTGCA-3’, mCox 1 forward

5’-GGGAATTTGTGAATGCCACC-3’, mCox 1 reverse

5’-GGGATAAGGTTGGACCGCA-3’, mCox 2 forward

5’-CAGACAACATAAACTGCGCCTT-3’, mCox 2 reverse

5’-GATACACCTCTCCACCAATGACC-3, mBLT1 forward

5’-GGCTGCAAACACTACATCTCC-3’, mBLT1 reverse

5’-TCAGGATGCTCCACACTACAA-3’,mBLT2forward

5’AGCCTTGGCTTTCTTCAGTTC-3’, mBLT2 reverse

5’-CCCTCGAAGAGTCGAGTAAGG-3’ and mTNFα forward

5’-TCTGTCTACTGAACTTCGGGGTGA-3’, mTNFα reverse

5’- GAAGTTGACGGACCCCAAAA-3’, IL-1β forward

5’- GCCTGCCTGAAGCTCTTGTT-3’, IL-1β reverse

Cycling conditions were: 50°C for 2 min, 95°C for 10 min, followed by 40 cycles of 95°C for 15 s and 60°C for 1 min. Melting curve analyses were performed for all real-time PCR runs to confirm amplification of only one product and to eliminate false positive results. mFFA4 and mFFA1 expression was defined relative to GAPDH and hFFA4 was presented relative to the reference gene ACTB and determined using the equation 2-dCq, where dCq = (Cq target gene - Cq ACTB RNA) and in some cases, this expression was arbitrarily multiplied by the factor 106 for clearer presentation. Expression of CXCL1 and TNFα was defined relative to HPRT using the 2^−ΔΔCt^ method.

### Staining of lung tissues for β-galactosidase activity

Wild type C57BL/6N or FFA4-KO(*β*gal) adult female mice (8-10 wk old) were sacrificed by cervical dislocation followed by exsanguination. Lung tissue was isolated and snap-frozen on OCT media. Frozen lung tissues were sectioned at 5 μm thickness on a cryostat and stained for β-galactosidase activity using GALS kit (Sigma) according to the manufacturer’s instructions.

### Immunostaining for the FFA4 receptor in the mouse lung

Mice were sacrificed by cervical dislocation, ensuring that the trachea remained intact. The chest cavity and trachea were exposed, and the trachea was cannulated. The lungs were gently overinflated with ice-cold 4% (wt/vol) paraformaldehyde (PFA/PBS). Following fixation, lungs were immediately removed and further fixed overnight in 4% (wt/vol) PFA/PBS at 4°C. Lungs were processed in paraffin wax, and sections from each lobe were taken using a microtome. Slices were either stained with haematoxylin and eosin or processed for antigen retrieval. Following antigen retrieval, sections were washed in Tris-buffered saline (TBS) plus 0.1% Triton X-100 (pH7.4) and blocked for 2 hr at RT in TBS, 0.1% Triton X-100, 10% (vol/vol) goat serum, and 1% (wt/vol) BSA. Sections were incubated with an in-house rabbit anti FFA4 receptor antiserum ^18^ (1:500 in blocking buffer) overnight at 4 °C. Sections were washed three times in TBS plus 0.1% Triton X-100 (pH7.4) and then incubated with fluorescent secondary antibodies (1:1000 in blocking buffer mouse Alexa Fluor 488 from Molecular Probes, Inc. and 1:500 in blocking buffer anti α-smooth muscle actin-Cy3 Ab from Sigma-Aldrich) for 2 hr at RT. Following three washes with TBS plus 0.1% Triton X-100 (pH7.4), slices were mounted in Vectorshield hardset mounting medium containing DAPI. ASM cells were seeded onto 8 well chamber slides and bronchial epithelial cells were cultured on 25-mm glass cover slips in 6 well plates to subconfluence. Cells were then fixed using methanol and blocked with PBS/3% BSA. Cells were incubated with a rabbit polyclonal anti-FFA4 (Novus Biologicals) verses rabbit isotype control (Immunostep, Salamanca, Spain), followed by anti-rabbit RPE conjugated secondary antibody (BioRad, Watford, UK). All images were taken using either a Zeiss Axiovert 200M microscope with a Colibri illumination system with Axiovision 4.8 software (Zeiss) or a Zeiss LSM 510 META NLO microscope with Zen 2009 software (Zeiss). Images were taken using the same magnification and microscope settings.

### Preparation of precision cut lung slices (PCLS)

Wild type female C57BL/6N or FFA4-KO(*β*gal) adult mice (8–10 wk old) were sacrificed by cervical dislocation followed by exsanguination. The trachea was exposed and inserted with a cannula. Lungs were inflated using 1.0 ml of 2% (wt/vol) low gelling agarose dissolved in HBSS buffer (supplemented with 10 mM Hepes, 1 mM MgCl_2_, 2 mM CaCl_2_ (pH 7.4)), followed by a 0.1-ml air bolus to force the agarose out of the airways and into the alveoli. The inflated lungs were dissected from the thoracic cavity and sliced using an automated vibrating microtome (VT1200S; Leica Biosystems) at a thickness of 200 μm. Lung slices were placed in ice-cold HBSS buffer supplemented with 10 mM Hepes, 1 mM MgCl_2_, 2 mM CaCl_2_ (pH 7.4) and then transferred to DMEM cell culture medium. Lung slices were placed in a humidified incubator (37°C, 5% CO_2_) and the medium was changed every 30 min to 1 hr for 6 hr to remove any stress factors that are released during the slicing. The slices were then cultured overnight in DMEM and used in experiments the following day.

### Airway contraction and relaxation in mouse PCLS

PCLS were placed in a 30 mm^2^ tissue culture dish in 3.0 mL of assay buffer (HBSS supplemented with 10 mM Hepes, 1 mM MgCl_2_, 2 mM CaCl_2_, pH 7.4). The airway was located using a microscope (Zeiss Axiovert 200M), and the slice was held in place using a stainless-steel ring. The camera was positioned so that a live video feed (AxioCAM) of the airway could be viewed. Media/DMSO were added for baseline measurements. A baseline image was taken, followed by the administration of the appropriate ligands. Images were collected every 1 min for 15-20 min per ligand addition. Airway luminal area was measured using the Axiovision software and data were plotted as a percentage of contractions of the initial (basal) airway size for the contractile agents. Log pEC_50_ and E_max_ values for each airway were derived from a concentration–response curve.

### Cell isolation and culture

Pure primary ASM bundles were isolated from bronchial biopsies and from lung resection material. ASM cells were cultured in DMEM with glutamax-1 supplemented with 10% FBS, penicillin (100U/ml), streptomycin (100µg/ml), amphotericin (0.25µg/ml), 100µM nonessential amino acids, and 1mM sodium pyruvate (Gibco). ASM was cultured and characterized for *α*-Smooth Muscle Actin (SMA) expression using a mouse monoclonal anti-SMA antibody (clone 1A4, Dako) or mouse IgG2a isotype control (clone DAK-G05, Dako) by flow cytometry and used between passage 2 to 6. Basal epithelial cells obtained from nasal brushings and bronchoscopy were grown on collagen (Advanced Bio Matrix, San Diego, California, USA) coated 12-well plates in bronchial epithelial growth medium (BEGM; Lonza, Berkshire, UK) including supplement SingleQuot BulletKit (Lonza), 0.3% Fungizone antimycotic (Invitrogen), and 1% antibiotic-antimycotic. The epithelial cells were expanded onto collagen-coated T75cm^2^ flasks and replenished with fresh medium three times a week.

### Detection of PGE_2_ release by ELISA

PCLS from the lungs of wild-type and FFA4 knock-out mice were placed in 24 well plates containing assay buffer and then treated with either vehicle or TUG-891 for 15 min at 37oC. Following treatment, the supernatant was taken and analysed using PGE2 ELISA from Abcam (ab133021) according to the manufacturer’s instructions. **BALF supernatants from mice undergoing in vivo experiments were also analysed using PGE2 ELISA to detect for the release of PGE2 into the alveolar spaces**. The amounts of PGE2 released from the **lung slices and BALF** was determined by interpolating the sample absorbance from the standard curve.

### Detection of LTB4 by ELISA

LTB4 concentrations in BALF of mice that had been chronically exposed to ozone and treated with TUG891 were analysed using LTB4 ELISA from R&D Systems (KGE006B) according to the manufacturer’s instructions. The amounts of LTB4 released into the BALF was determined by interpolating the sample absorbance from the standard curve.

### Ca^2+^ signalling in lung slices

PCLS from wild type or FFA4-KO(*β*gal) mice were washed twice in assay buffer (HBSS supplemented with 10 mM Hepes, 1 mM MgCl_2_, 2 mM CaCl_2_ (pH 7.4)) and incubated in the buffer containing 4 μM Fura-2:00 AM (Molecular Probes), 1 mg/mL BSA, and 2.5 mM probenecid for 30 min at 37 °C. After loading, the slice was placed in a prewarmed chamber containing assay buffer. The slice was held in place using a stainless-steel ring and measurements were made on a Zeiss Axiovert 200 inverted epifluorescence microscope with a 40× oil-immersion objective. Loaded lung slices were excited at 340 nm and 380 nm at a sample rate of 0.67 Hz by means of an excitation wheel. Sequential fluorescent image pairs were recorded at wavelengths >510 nm via a cooled ORCA-ER CCD camera (Hamamatsu Photonics) and processed with MetaFluor software (Molecular Devices). Free intracellular Ca^2+^ signal was expressed at a ratio of 340:380 or change of the ratio of fluorescence.

### Airway hyper-responsiveness (AHR) assessment associated with ozone exposure models

Mice were anesthetized with an intraperitoneal injection of anaesthetic solution containing ketamine and xylazine diluted in physiological saline. Mice were tracheostomised and ventilated (MiniVent type 845, EMMS, Hants, UK) rate: 250 breaths/min and tidal volume: 250 ml) and resistance/compliance measurements were made in a whole-body plethysmograph with a pneumotachograph connected to a transducer (EMMS, Hants, UK). Transpulmonary pressure was assessed via an oesophageal catheter (EMMS, Hants, UK). Instantaneous calculation of pulmonary resistance (RL) was obtained. FFA4 receptor ligands were administered with an AeronebH Lab Micropump Nebulizer (EMMS, Hants, UK), and RL was recorded for a 3-min period followed by administration of acetylcholine (128 mg/ml) (Sigma, Dorset, UK). RL after each ligand addition was expressed as percentage change from baseline RL measured following nebulized PBS (Sigma, Dorset, UK). The change in RL caused by acetylcholine in the presence and absence of FFA4 receptor ligands relative to basal RL was taken as a measure of airway responsiveness.

### Assessment of cigarette mouse model lung function

Mice were anesthetized (50 μl/10 g i.p.) with a mixture of xylazine (2 mg/ml, Troy Laboratories) and ketamine (40 mg/ml, Ceva). Mice were cannulated *via* the trachea and ventilated with a tidal volume of 8 ml/kg at a rate of 450 breaths/min. FlexiVent^TM^ apparatus (FX1 System; SCIREQ, Montreal, Canada) was used to assess airway resistance as previously described ^63–65^. Mice were ventilated for 5 mins before baseline recordings were obtained using single frequency (2.5Hz, 1.2s long; Snapshot-150) and broadband low frequency (1-20.5 Hz, 8s long; Quick-Prime-8) forced oscillation manoeuvres. Mice were then challenged with aerosolised PBS for 1 min and Snapshot-150 and Quick-Prime-8 recorded, followed by aerosolised TUG-1197 (0.364 mg/ml) for 3 mins (Snapshot-150 and Quick-Prime-8 recorded), and then aerosolised methacholine (30 mg/ml, Sigma) for 3 mins (Snapshot-150 and Quick-Prime-8 recorded). Rrs (from Snapshot-150) and Rn (from Quick-Prime-8) were analysed as a percentage change from TUG-1197.

### Immunohistochemistry of human lung biopsies

Bronchial biopsy specimens from healthy subjects were processed and embedded as described in^66^. 2 micron sections were then processed on the Autostainer Link 48 platform (Dako AS480) using an automated staining protocol for the FFA4 antiserum (4 µg/ml) and its rabbit polyclonal isotype control (Immunostep, Salamanca, Spain) using EnVision FLEX kit (Dako). Pictures were obtained using a Carl Zeiss Imager Z2 microscope and AxioCam HRc digital camera (Carl Zeiss, Germany).

### Ex vivo airway contraction and relaxation in human PCLS

Experiments were performed by Biopta Ltd (Glasgow, UK). In brief, quaternary branches of human bronchi from 3 healthy donors were dissected free from the surrounding parenchyma. Bronchi were cut into 2 mm rings and mounted in 5 ml myographs (Danish Myo Technology A/S) containing physiological saline solution (composition: 119.0 mM NaCl, 4.7 mM KCl, 1.2 mM MgS04, 24.9 mM NaHCO3, 1.2 mM KH2PO4, 2.5 mM CaCl2, and 11.1 mM glucose), aerated with 95% O2 / 5% CO_2_ gas mix, warmed and maintained at approximately 37°C. The tissue was held at a resting tension of 1-1.5 g, and the buffer was exchanged every 15 min for 2 h with a continuous digital recording of muscle force. Following this equilibration period, a pre-determined EC_80_ concentration of acetylcholine (10 μM) was added to the chamber for 5 min or until the plateau was reached. Concentration-response curves of TUG-891 were constructed (10 nM–100 μM) with each concentration of the compound added to the tissue for 5 min or until the plateau was reached. At the end of the experiment, theophylline (2.2 mM) was added to induce maximal relaxation of the airways.

Studies of potential off-target effects of FFA4 agonists on prostanoid receptors Pharmacological profiling and determination of potential off-target effects of FFA4 agonists on prostanoid receptors were conducted (TUG-1197, study ID, FR095-0002695, TUG-891, study ID 100025966) by Eurofins/Cerep Panlabs (Celle-Lévescault France), a contract research organisation specialising in providing pharmacological screening services. TUG-891 and TUG-1197 were screened at a 10 μM concentration in duplicate for potential to bind to and activate or inhibit a panel of prostanoid receptors, EP_1-4_. Details and experimental conditions for these assays can be found online at https://www.eurofinsdiscoveryservices.com/cms/cms-content/services/in-vitro-assays/gpcrs/.

### Assessment of airway smooth muscle contraction by collagen gel analysis

Contractile properties of human airway smooth muscle (HASM) cells were assessed using a collagen gel contraction assay. Collagen gels were impregnated with HASM cells (2.5 × 10^5^ cells) resuspended in DMEM with GlutaMAX-1 supplemented with penicillin (100 U·mL^−1^), streptomycin (100 μg·mL^−1^), amphotericin (0.25 μg·mL^−1^), non-essential amino acids (100 μM) (Invitrogen, Paisley, United Kingdom) and sodium pyruvate (1 mM). Next, 500 μL of gel mixture was added to each well of a PBS 2% BSA pre-coated 24-well plate and allowed to polymerize at 37°C for 90 min. After polymerization, 500 μL DMEM with GlutaMAX-1 (supplemented as above) was added to each well, and the gel was detached from the plastic surface to allow free contraction. Carbachol at a concentration of 100µM was added to induce contraction where appropriate for 1 hour prior to the addition of vehicle control, TUG-891 or AH-7614. Images were captured over 3 hours and at 24 hours. Gel area was measured as a percentage of control basal well area using ImageJ software. All gel conditions were performed in duplicate.

### ERK1/2 phosphorylation assay

mFFA4 signaling to phospho-ERK1/2 was detected by using the HTRF phospho-ERK1/2 (Thr202/Tyr 204) kit (Cisbio Biassays, Codolet, France). In brief, cells were seeded in 96-well plates at 30,000 cells/well and grown for 48 hours before being serum starved overnight at 37°C. Cells were stimulated with an agonist and lysed with 50 µl lysis buffer for 1 hour at room temperature. For concentration-response curves, cells were stimulated with increasing concentration of agonists for 5 minutes and for time-course experiments, cells were stimulated with maximal concentration of ligand (1 μM) for the indicated time. Lysates [16 µl] were transferred to 384-well plates and 4 µl premixed d2/cryptate antibodies [1:1 (v/v)] were added to each lysate. The reaction mixtures were incubated for 2 hours at room temperature and the plate was read on the ClarioStar plate reader (BMG Labtech).

### COX-2 inhibition in chronic ozone model

Mice (female C57BL6, age 10 - 12wk) were administered daily with either vehicle (1.5% DMSO), TUG891 alone (100 μM) or TUG891 in combination with the COX2 inhibitor celecoxib (50 μM), via intranasal delivery (25 μl volume). Mice were then exposed to 3 ppm ozone for 3 hr, twice a week for total duration of 3 weeks. Control animals exposed to normal air and vehicle treatment (1.5% DMSO) were included in the experiments. Following treatment, mice were humanely sacrificed via an overdose of pentobarbitone. Once death is confirmed, mice were tracheostomised and bronchoalveolar lavage fluids (BALFs) were obtained. Left lung lobe from each mouse was dissected and snap frozen for total RNA isolation and RT-qPCR.

### Detection of TNFα release by ELISA

BALF supernatants of mice from the in vivo COX-2 inhibition experiments were analysed using TNFα ELISA from Life Technologies (cat# # BMS6002) according to the manufacturer’s instructions. The amounts of TNFα released into the alveolar spaces were determined by interpolating the sample absorbance from the standard curve.

## Statistical analysis

Data were analysed by ANOVA with Bonferroni post-hoc and Student’s t-test as indicated.

## Supplementary Data

**Fig. S1.**
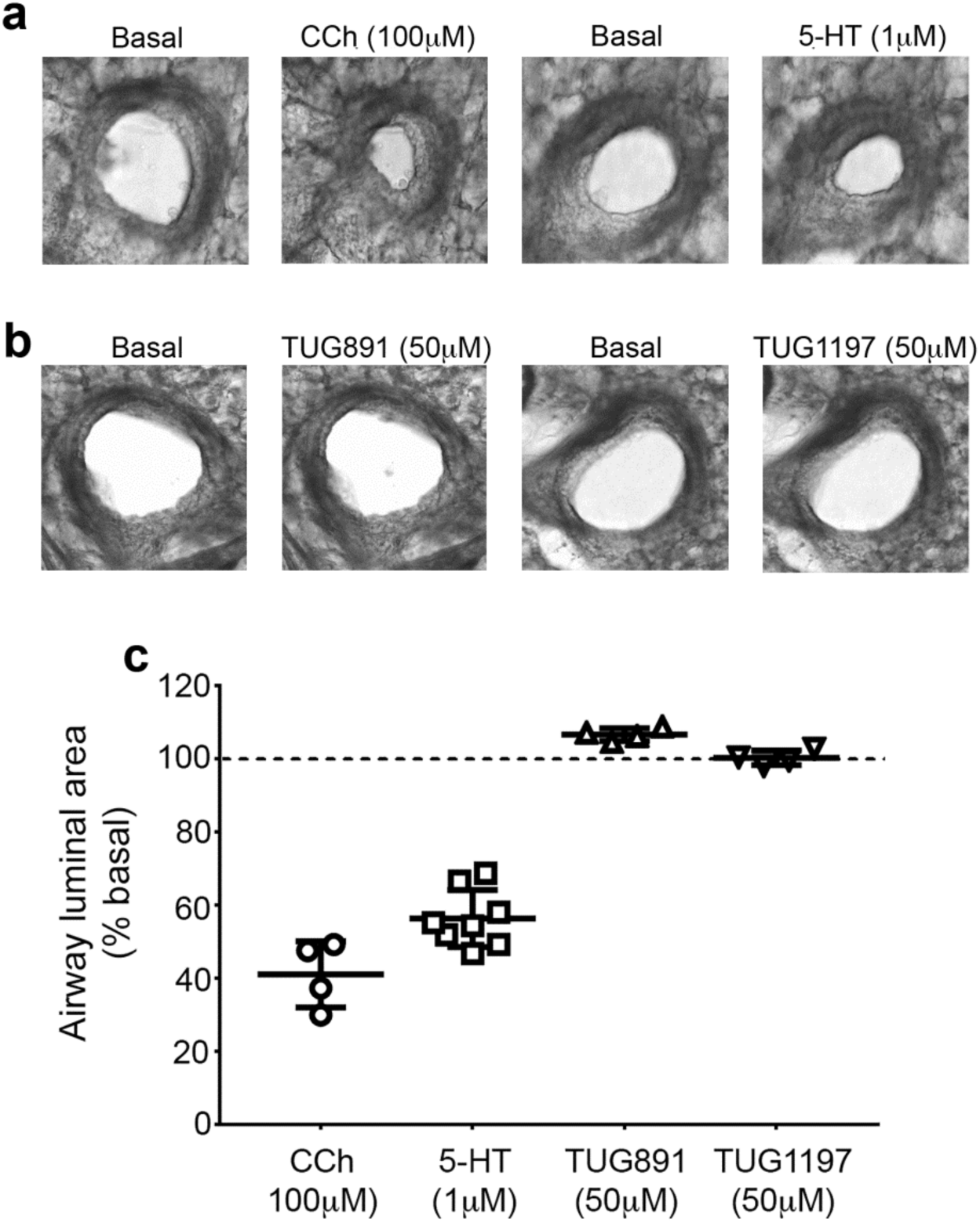
FFA4 activation does not cause airway contraction in murine PCLS. (**a**) Representative images of airway responses evoked by **(a)** the muscarinic receptor agonist carbachol (CCh) and serotonin (5-HT) and (**b**) by the FFA4 agonists TUG-891 and TUG-1197. **(c)** Quantification of the data shown in **a** and **b** from at least four animals. The data are the mean ± S.D.

**Fig S2.**
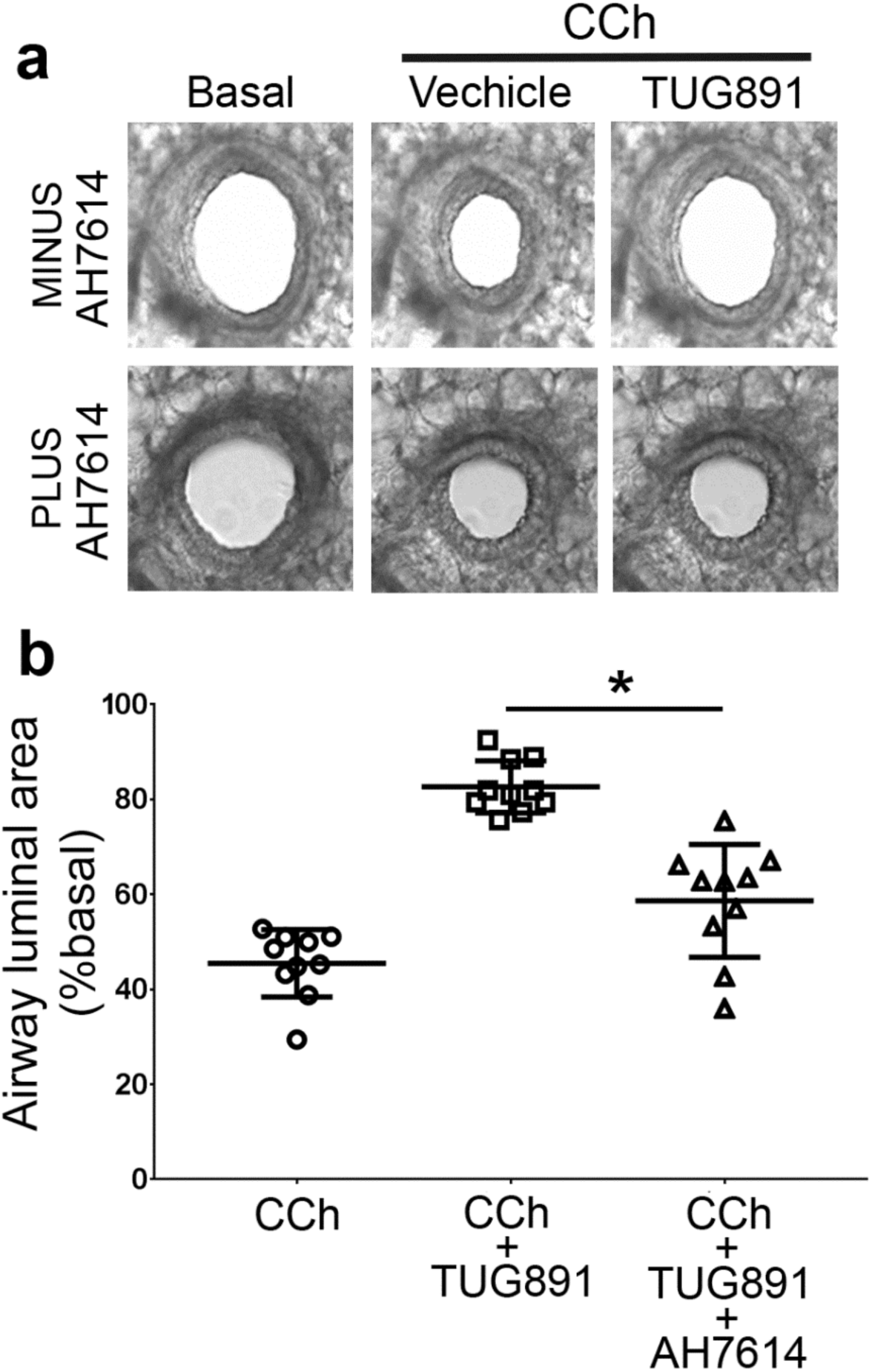
TUG-891 mediated relaxation in pre-contracted PCLS was reduced by the FFA4 antagonist AH7614. **(a)** Representative images of the effect of the FFA4 antagonist AH7614 on the TUG891-mediated relaxation response in PCLS pre-contracted with carbachol (CCh 100μM). **(b)** Quantification of the experiment in **a** from at least ten animals where *p<0.05 as determined by ANOVA with Bonferroni post-hoc test.

**Fig S3.**
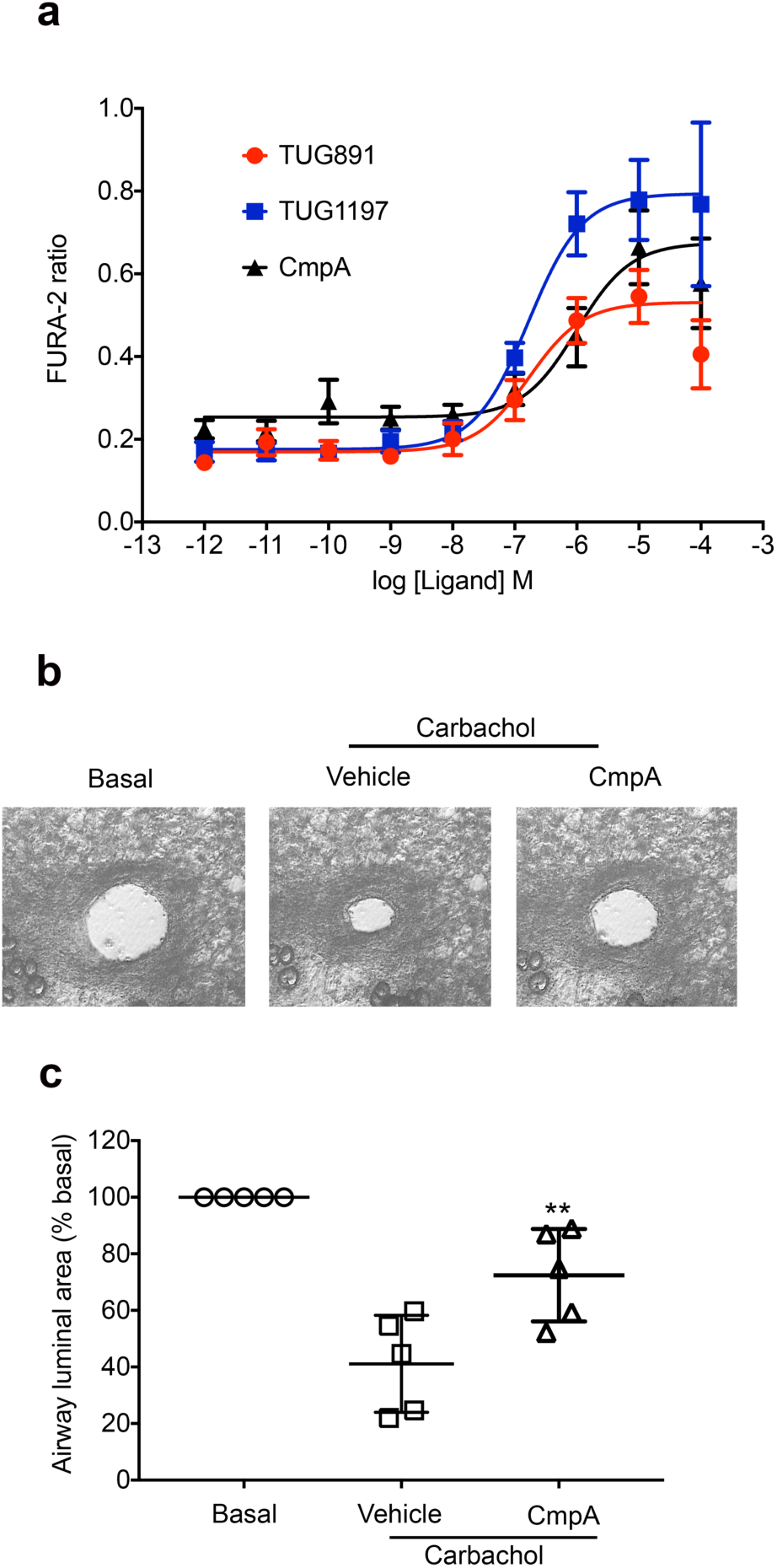
FFA4 agonist Compound A mediates ASM relaxation in pre-contracted PCLS. **(a)** Calcium concentration response curves in CHO Flp-In cells expressing mouse FFA4 stimulated with TUG-891, TUG-1197 or Compound A (CmpA). **(b)** Representative images of the effect of Compound A (CmpA, 50μM), on PCLS pre-contracted with carbachol (CCh 100μM). **(c)** Quantification of the experiment in **b** where significant difference between vehicle and compound A was determined by ANOVA with Bonferroni post-hoc test. **p<0.01.

**Fig S4.**
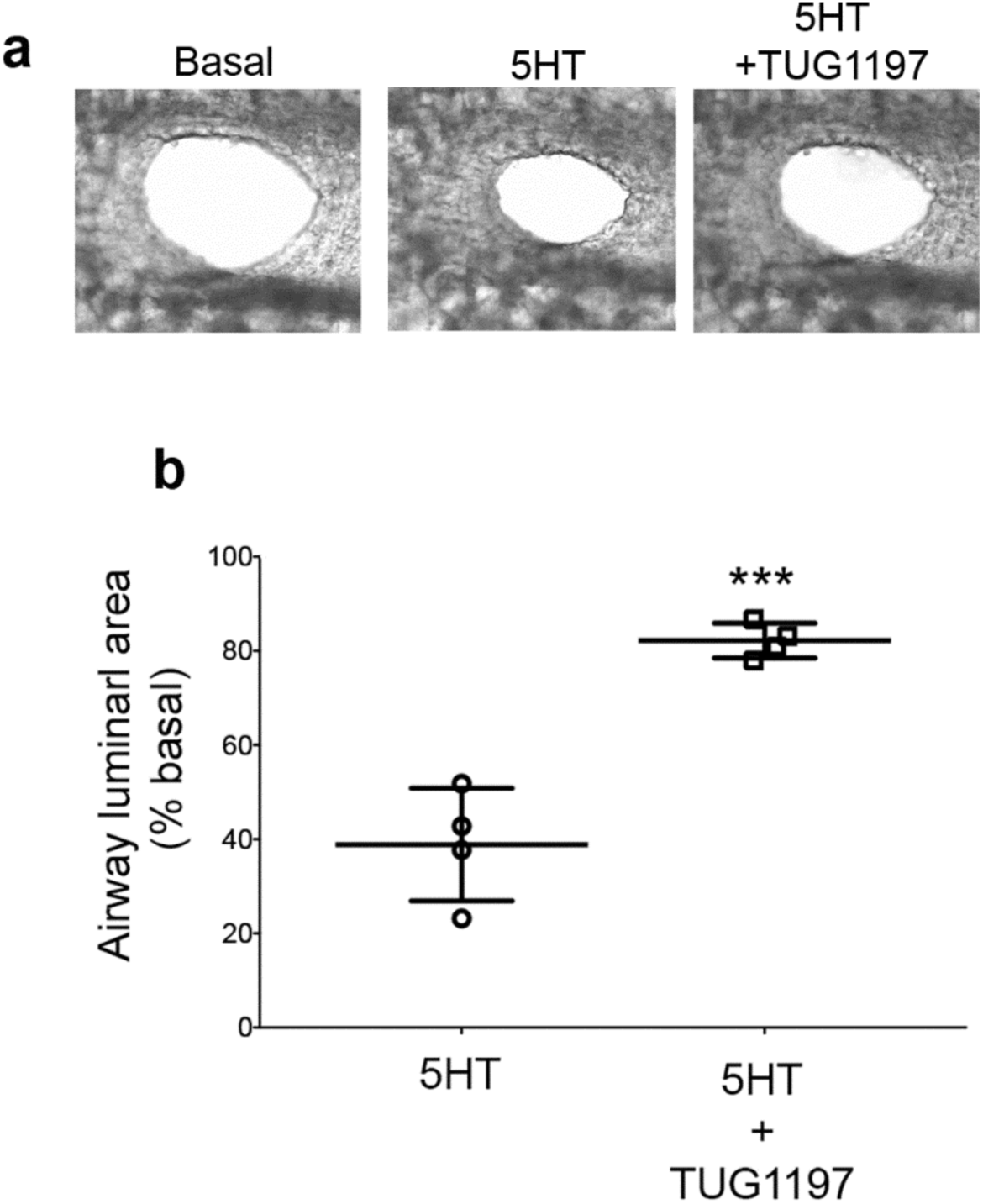
FFA4 agonism mediates relaxation in airways pre-contracted with serotonin. (**a**) Representative images of PCLS pre-contracted with serotonin (1 μM) followed by treatment with TUG-1197 (50μM). **(b)** Quantification of images presented in panel a. Data are mean ± S.D. of at least four animals where ***p<0.005 as determine by Student’s t test.

**Fig S5.**
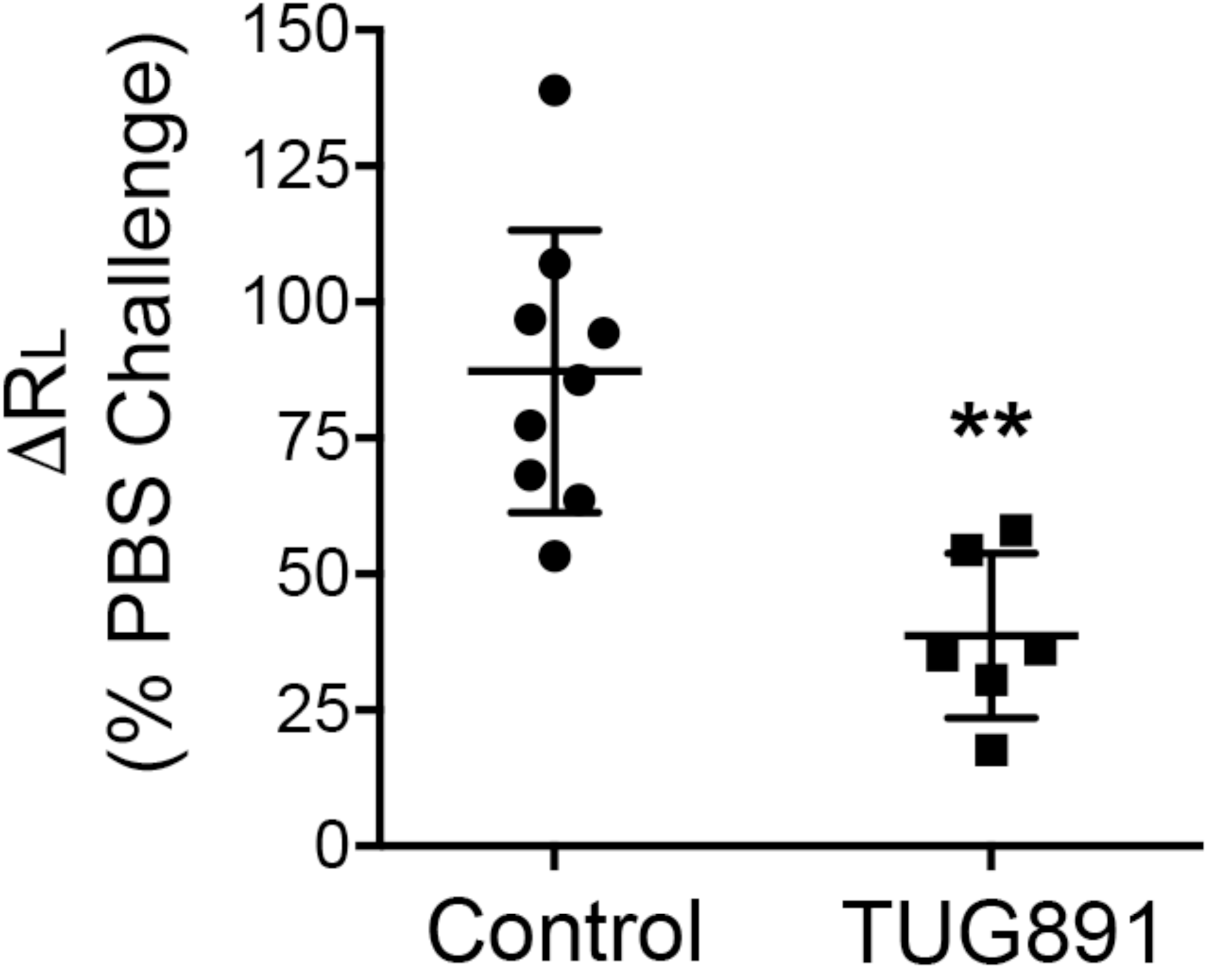
FFA4 agonist TUG-891 reduces airway resistance in wild type C57BL/6 mice. Vehicle (Control) or TUG-891 (0.356 mg/ml) were nebulized with acetylcholine and administered to wild type mice and lung resistance measured. The data is expressed as the mean ± S.D. The data shown were analysed by unpaired t test where **p<0.01.

**Fig S6.**
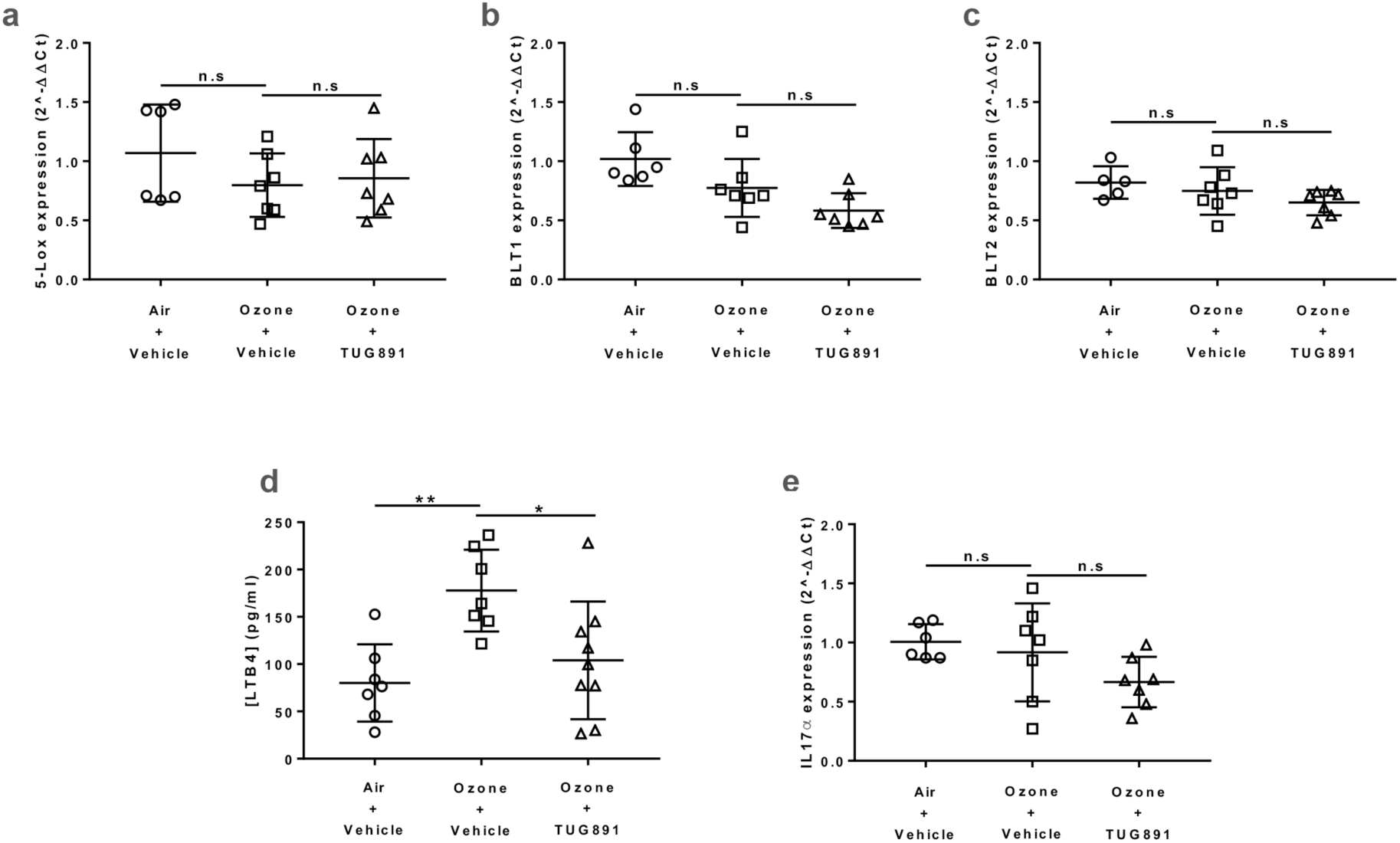
FFA4 agonism did not significantly alter transcript levels of 5-Lox, BLT1, BLT2 and IL17*α* but increased expression of LTB4 in chronic ozone model. FFA4 agonism did not significantly alter transcript levels of 5-Lox, BLT1, BLT2 and IL17*α* but significantly reduced LTB4 proteins in chronic ozone model. Mice were exposed to either control air or 3ppm ozone for 3hrs, twice a 3 week for a total duration of 3 weeks. Prior to each ozone exposure, mice were administered with either vehicle or TUG891 (0.036 mg/ml). Mice were then sacrificed and RT-qPCR analysis was performed in lung tissue to identify changes in the transcript levels of **(a)** 5-Lox, **(b)** BLT1, **(c)** BLT2 and **(e)** IL17a genes. ELISA assay was performed in BALF to identify changes in LTB4 protein level **(d).** Data represent the means ± S.D. of 6-9 animals.

**Fig S7.**
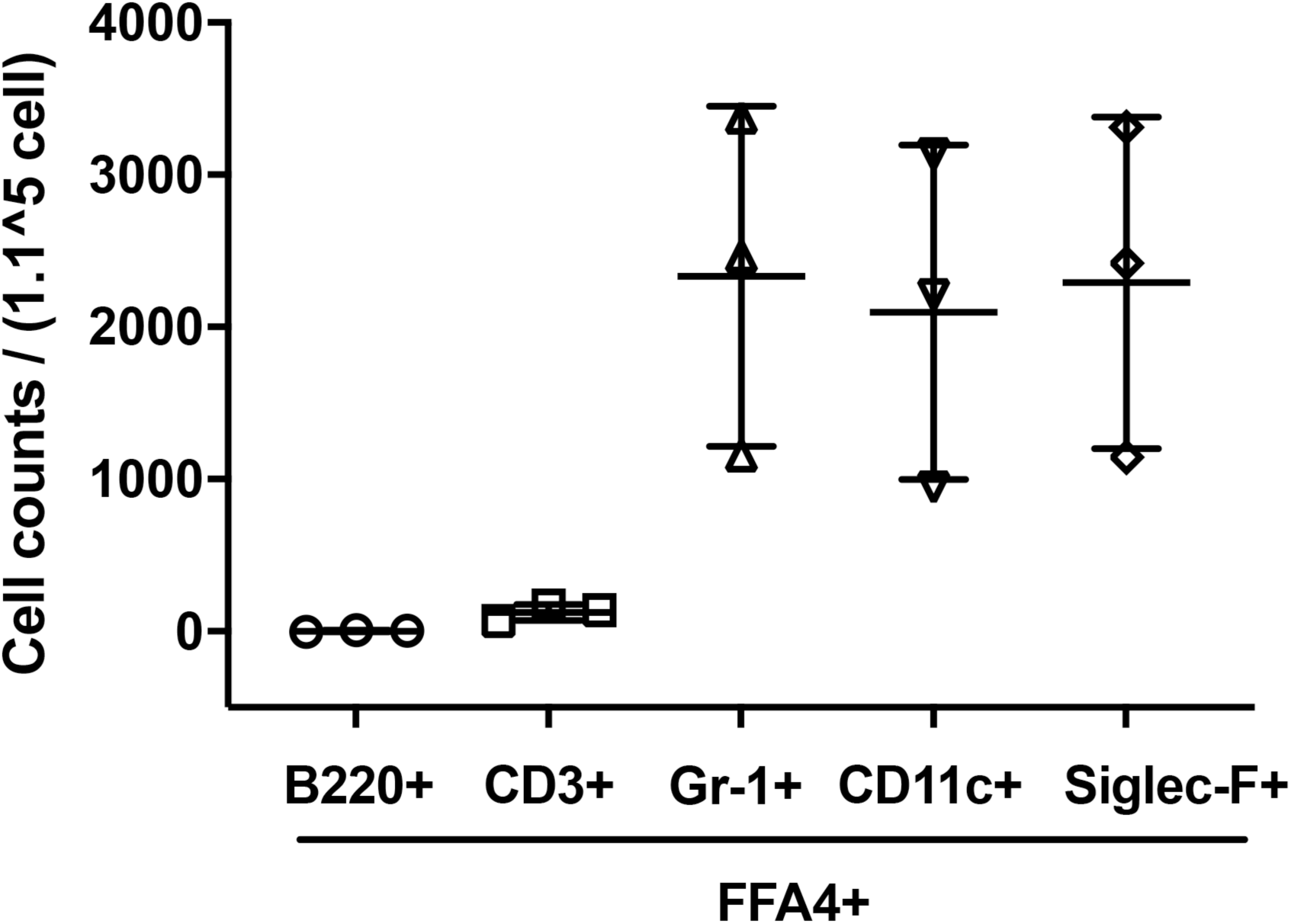
FFA4 is expressed in Gr-1+, CD11c+, and siglec-F+ immune cells. Mice were exposed to ozone for 3hrs, twice a 3 week for a total duration of 3 weeks. Mice were sacrificed and flow cytometry analysis of BALF cells was performed to detect for co-staining of FFA4 with common markers (B220, CD3, Gr-1, CD11c and siglec-F) of immune cells. Data represent the means ± S.D. of 3 animals.

**Fig S8.**
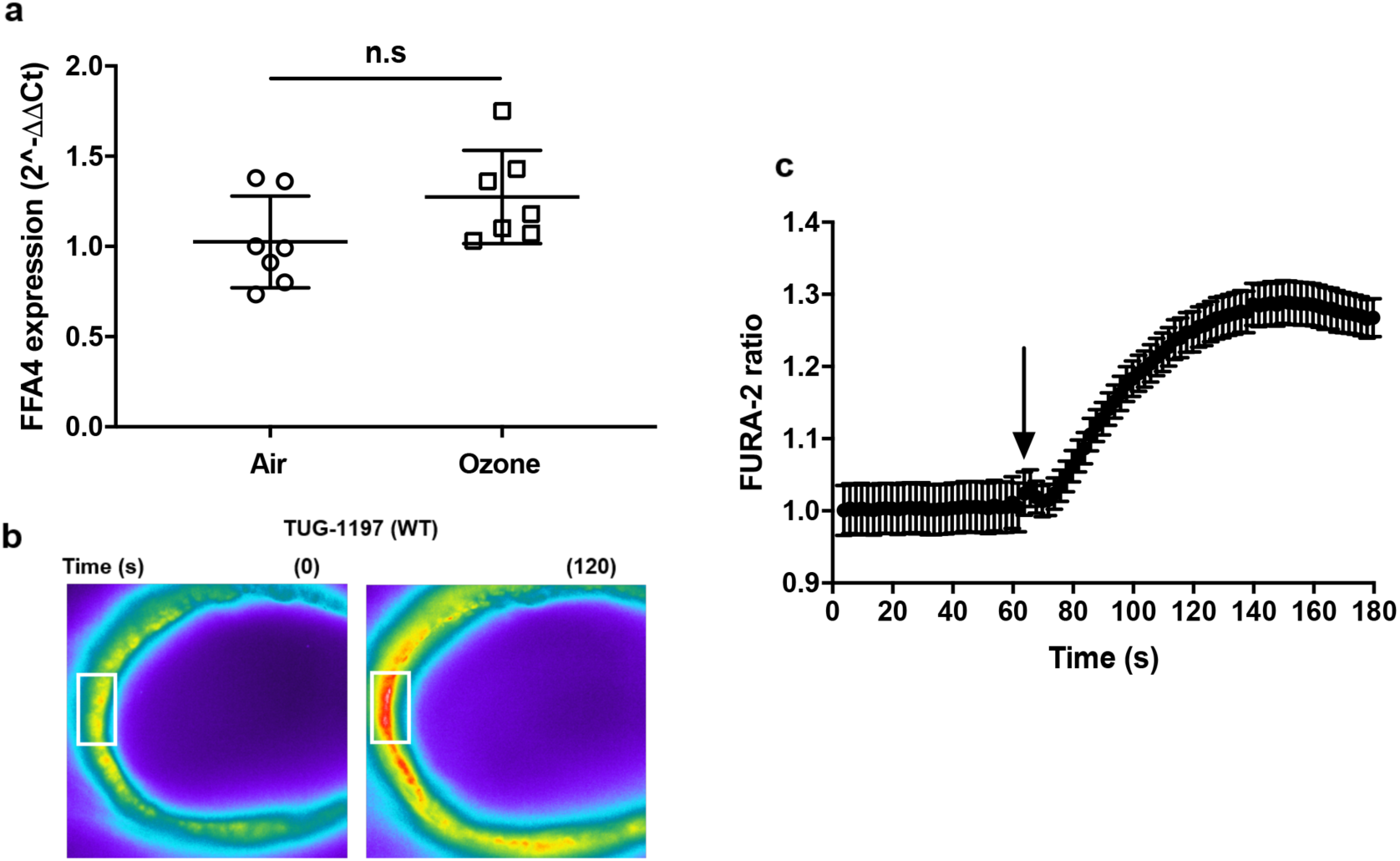
FFA4 is expressed normally and fully functional in ozone-exposed mice. Mice were exposed to either control air or 3ppm ozone for 3hrs, twice a 3 week for a total duration of 3 weeks. Mice were sacrificed and RT-qPCR analysis was performed to identify changes in the expression of **(a)** FFA4 gene. PCLS were prepared from ozone-exposed mice and stimulated with TUG891 to detect for calcium mobilisation **(b-c).** Data represent the means ± S.D. of 4-7 animals.

**Fig S9.**
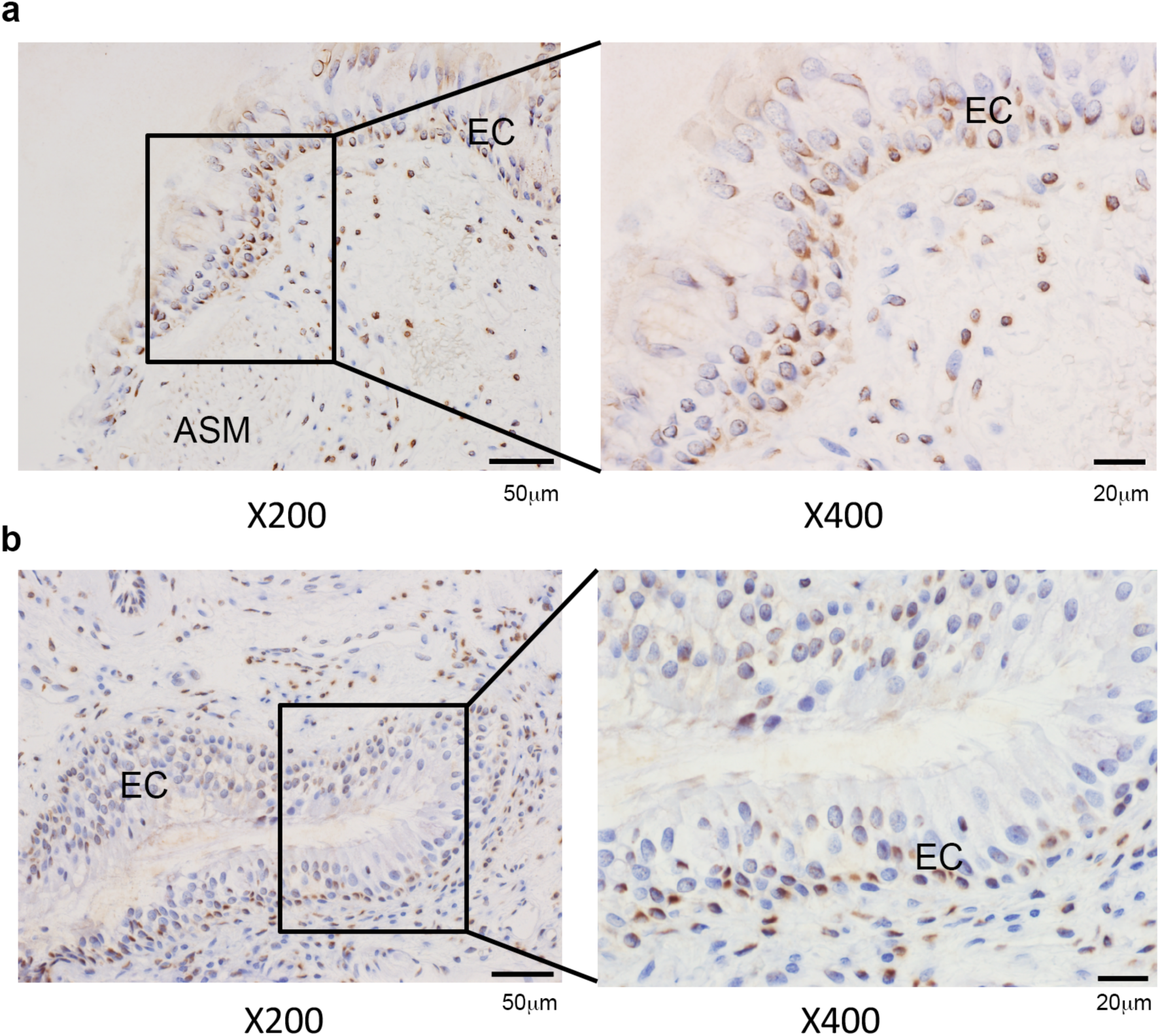
Expression of FFA4 in human lung. **a,b** Two representative photomicrographs of normal human bronchial biopsy sections stained with the human FFA4 selective antiserum (4 µg/ml). The boxes represent the zoomed regions. EC = epithelial cell layer, ASM = airway smooth muscle. LP = Lamina propria.

**Fig S10.**
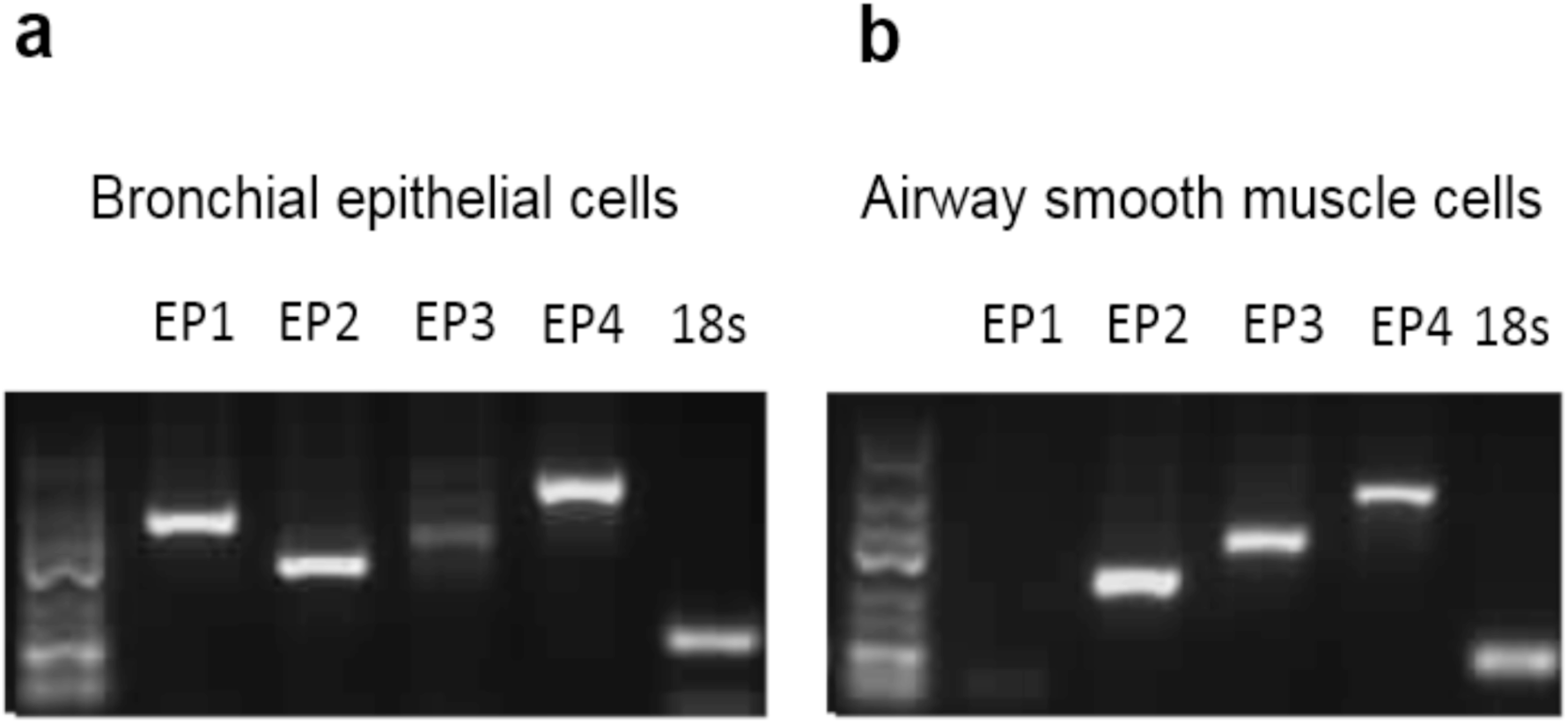
PGE2 receptors are expressed in human bronchial epithelial cells and human airway smooth muscle cells. Transcript levels of EP1, EP2, EP3, EP4 and 18s (as a control) were determined in **(a)** bronchial epithelial cells and **(b)** ASM cells using RT-PCR. Shown is a representative gel of 5 donors. The expected sizes of PCR products are as follows; EP1= 324bp, EP2=216bp, EP3=300bp, E4=434bp and 18s=93bp.

**Fig S11.**
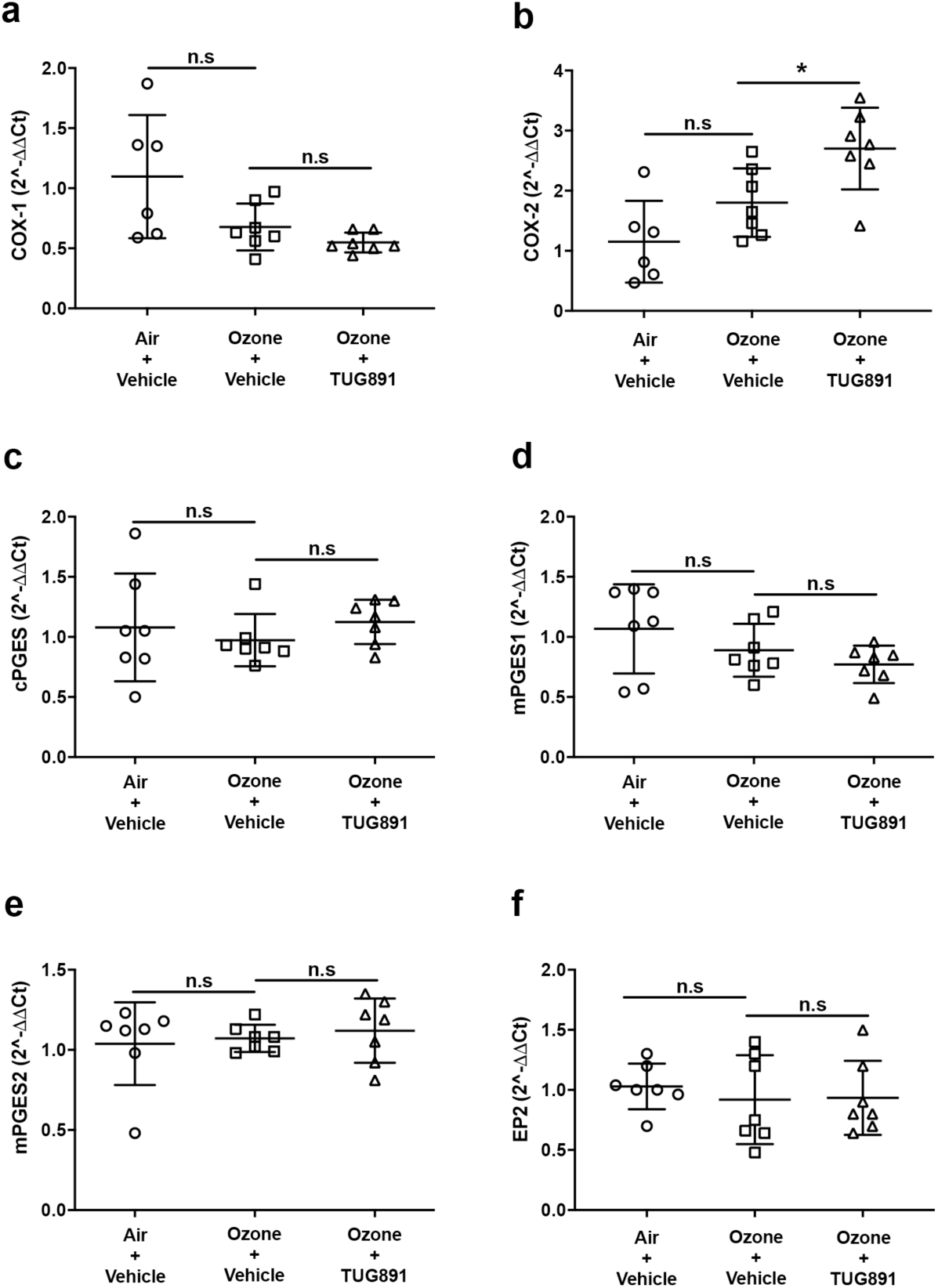
FFA4 agonism upregulates the transcript levels of COX-2 in chronic ozone model while the transcript levels of other genes involved in the biosynthesis and biological actions of PGE2 remained unaltered. Mice were exposed to either control air or 3ppm ozone for 3hrs, twice a 3 week for a total duration of 3 weeks. Prior to each ozone exposure, mice were administered with either vehicle or TUG891 (0.036 mg/ml). Mice were then sacrificed and RT-qPCR analysis was performed to identify changes in the transcript levels of (a) COX-1, (b) COX-2, (c) cPGES, (d) mPGES1, **(e)** mPGES2 and **(f)** EP2. Data represent the means ± S.D. of 6-7 animals.

**Fig S12.**
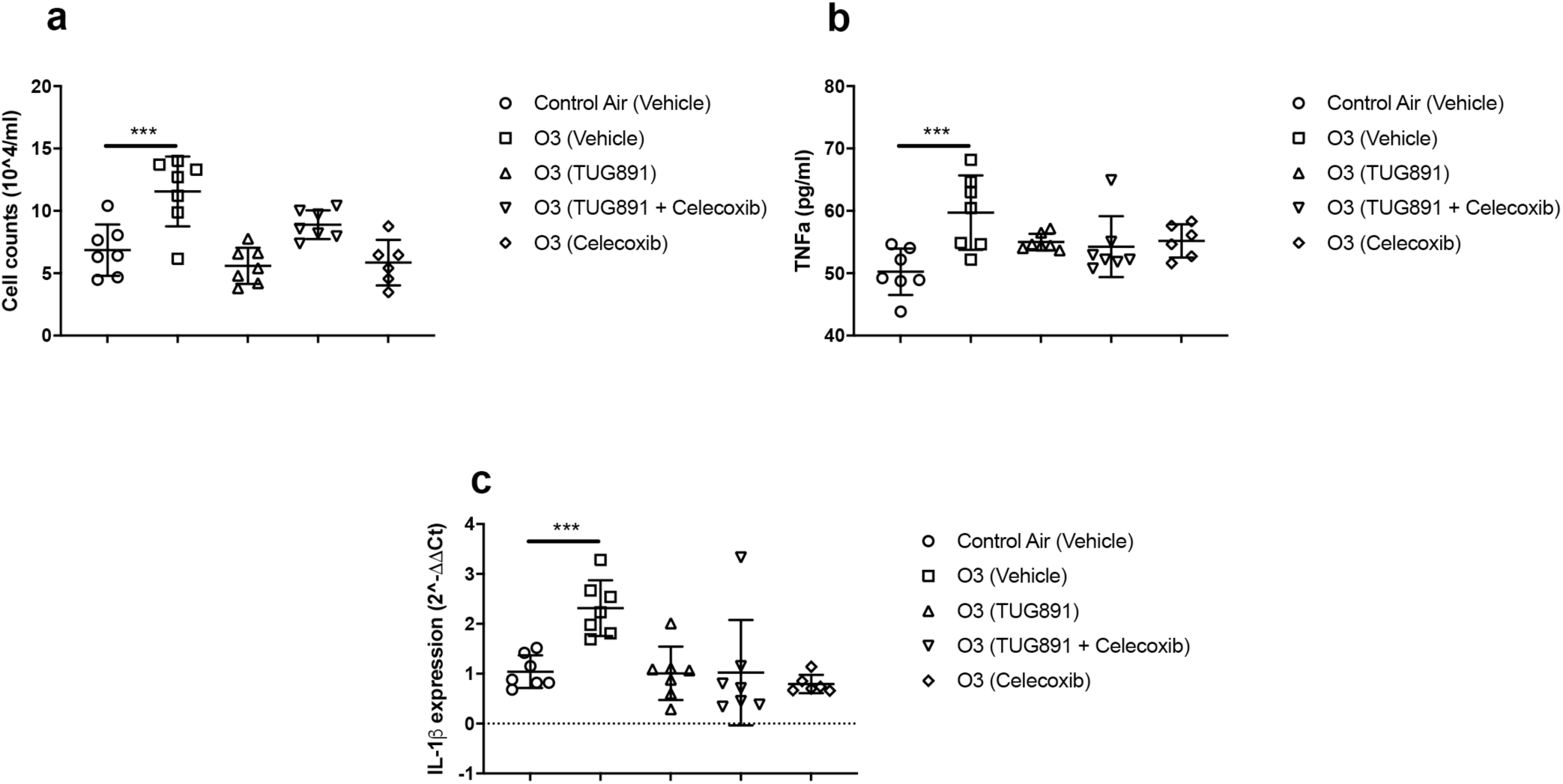
FFA4 activation mimics the anti-inflammatory effects of a COX-2 inhibitor but is not altered by the presence of COX-2 inhibition. (a) Total cell counts and (b) TNFα levels in the BALFs and (c) IL-1β expression in the lungs of mice exposed to normal air or chronically exposed to ozone treated with either vehicle (1.5% DMSO), TUG891 alone (100 μM), TUG891 in combination with the COX-2 inhibitor, Celecoxib (50 μM) or Celecoxib alone. Data represent the means ± S.D. of at least 6 animals where ***p<0.001, as determined by ANOVA with Bonferroni post-hoc test.

**Table S1.** FFA4 agonists do not act as off-target agonists at prostanoid receptors. The effect of 10 μM of either TUG-891 or TUG-1197 to stimulate each of the EP1-4 prostanoid receptors was compared to the maximal effect of a reference agonist. Data are presented as mean +/-range from two independent experiments.

| Receptor | Ligand | Response<br>(% control agonist) |
| --- | --- | --- |
| EP<sub>1</sub> | TUG-891 | 0.9 +/- 0.0 |
| EP<sub>1</sub> | TUG-1197 | 2.1 +/- 2.1 |
| EP<sub>2</sub> | TUG-891 | 1.2 +/- 0.4 |
| EP<sub>2</sub> | TUG-1197 | -3.8 +/- 1.0 |
| EP<sub>3</sub> | TUG-891 | -8.4 +/- 4.2 |
| EP<sub>3</sub> | TUG-1197 | -1.2 +/- 0.7 |
| EP<sub>4</sub> | TUG-891 | 3.0 +/- 2.3 |
| EP<sub>4</sub> | TUG-1197 | 0.0 +/- 1.9 |

**Table S2.**
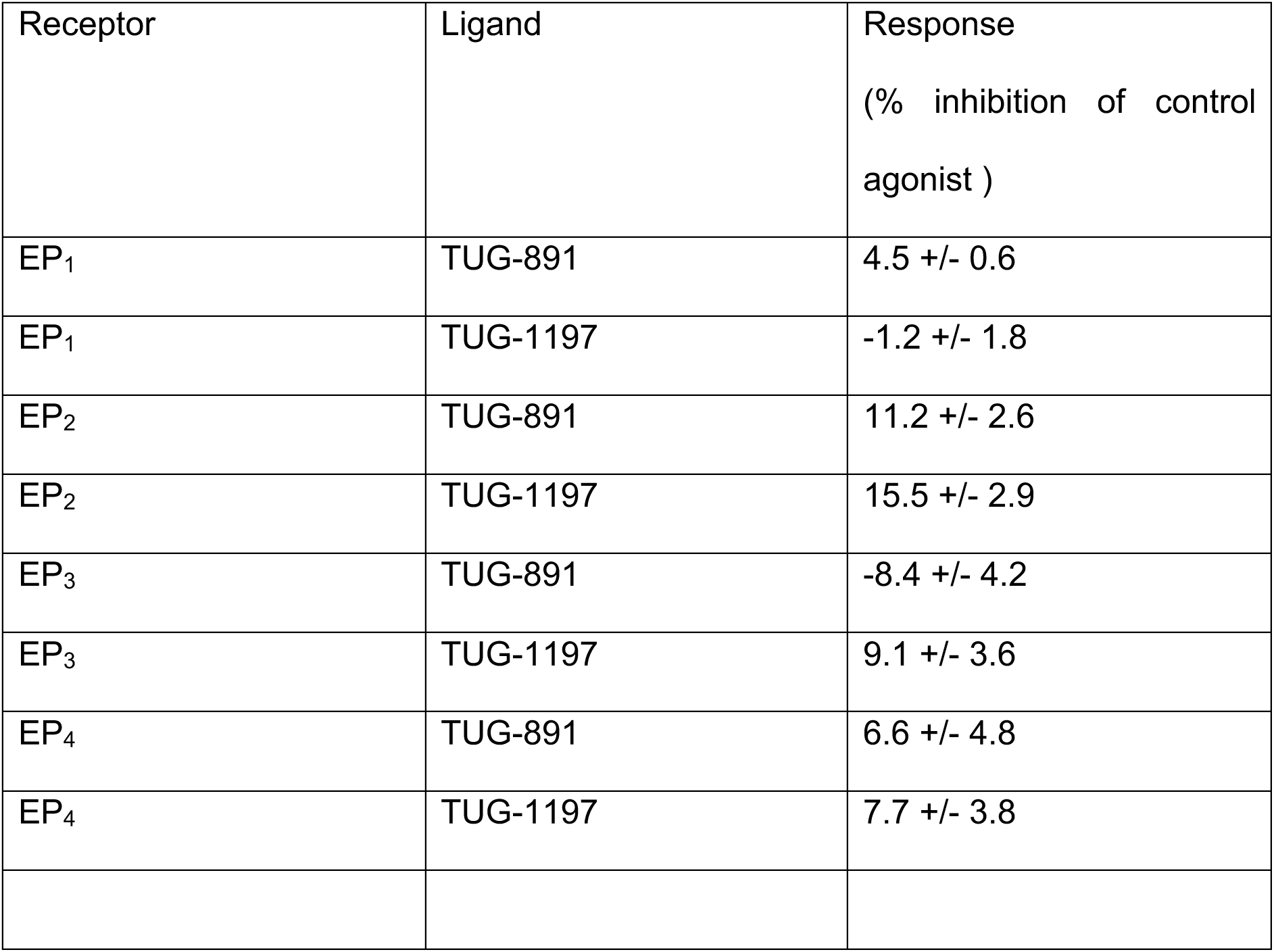
FFA4 agonists do not act as off-target antagonists at prostanoid receptors. The effect of 10 μM of either TUG-891 or TUG-1197 to inhibit response to a control agonist at each of the EP1-4 prostanoid receptors was assessed. Data are presented as mean +/-range from two independent experiments.

## Funding

This study is funded by the following Biotechnology and Biosciences Research Council (BBSRC) grants BB/K019864/1 (GM) and BB/K019856/1 (ABT) and by the Medical Research Council (MRC), grant MR/R00305X/1 (GM and ATB). Funding for CHW and KFC from MRC grant G1001367/1 and for TU The Danish Council for Strategic Research (grant 11-116196). Funding and fellowships for CD and PMH from the NHMRC 1079187, 1059238 and Rainbow foundation. PMH was also funded by a Fellowship and grants from the NHMRC (1079187, 1175134). RP received funding from The Royal Society and Welcome Trust Institutional Strategic Support Fund. This paper also contains independent research funded by the National Institute for Health Research (NIHR) Leicester Respiratory Biomedical Centre. The views expressed are those of the authors and not necessarily those of the NHS, the NIHR or the Department of Health.

## Acknowledgements

We thank Dr Evi Kostenis (Department of Pharmacy, University of Bonn, 53115 Bonn, Germany) for FR900359 and both Dr Carlos Azevedo and Dr Bharat Shimpukade for assistance with compound synthesis. We acknowledge the BSU facilities at the Cancer Research UK Beatson Institute (C596/A17196), the Biological Services and Flow Core at the University of Glasgow. We thank Maria Gaellman and Anne Ryan for support of the Tobin and Milligan laboratories.

## Author contributions

ABT,GM; Conceived and led the project. RP; Designed and coordinated the study. CEB; Designed and led the human airway studies. CD, PMH, RP, RK; Designed and performed the cigarette study. RP,KFC,CHW,JL; Contributed to the design and performed the ozone experiments. EAC,ZD,AGMA,EE; Performed pharmacology study. LC,CC; Performed human airway study. TU; generated FFA1/FFA4 ligands.

## Competing Interests

The authors declare no conflicts of interest.

## Data and materials availability

All reagents and materials used in this study may either be purchased or obtained from the corresponding authors.

## Notes

### Competing Interest Statement

The authors have declared no competing interest.

